# Cryo-EM analysis of complement C3 reveals a reversible major opening of the macroglobulin ring

**DOI:** 10.1101/2024.04.15.589532

**Authors:** Trine Amalie Fogh Gadeberg, Martin Høgholm Jørgensen, Heidi Gytz Olesen, Josefine Lorentzen, Seandean Lykke Harwood, Ana Viana Almeida, Marlene Uglebjerg Fruergaard, Rasmus Kjeldsen Jensen, Philipp Kanis, Henrik Pedersen, Emil Tranchant, Steen Vang Petersen, Ida Buch Thøgersen, Birthe Brandt Kragelund, Joseph Anthony Lyons, Jan Johannes Enghild, Gregers Rom Andersen

## Abstract

The C3 protein is the central molecule within the complement system and undergoes pattern-recognition-dependent proteolytic activation to C3b in the presence of pathogens and damage-associated patterns. Spontaneous pattern-independent activation of C3 occurs via hydrolysis, resulting in C3(H_2_O). However, the structural details of C3 hydrolysis remain elusive. Here, we show that the conformation of the C3(H_2_O) analog, C3MA, in which the C3 thioester is broken by aminolysis is indistinguishable from C3b except for the 77-residue anaphylatoxin (ANA) domain. In contrast, the reaction intermediate C3* formed during C3 adopts a dynamic conformation dramatically different from both C3 and C3MA/C3b. In C3*, unlocking of the macroglobulin (MG) 3 domain creates a large opening in the MG-ring through which the ANA domain translocates. In support of this mechanism, C3MA formation is inhibited by an MG3/MG4-interface-specific nanobody and prevented by linking the ANA domain to the C3 β-chain. Our study reveals an unexpected dynamic behavior of C3 where an exceptional conformational change allows the translocation of an entire domain through a large dynamic opening. These results form the basis for elucidation of the *in vivo* contribution of C3 hydrolysis to complement activation and offer a rational approach for modulation of C3(H_2_O) with the potential for preventing complement activation caused by intravascular hemolysis and surface contacts.

## Introduction

The complement system forms a first line of defense against pathogens and supports the removal of debris to maintain homeostasis in the circulation and tissues. Pattern-dependent activation of complement leads to cleavage of complement C3, where the anaphylatoxin (ANA) domain is released as C3a, and C3b is left for covalent deposition via its internal thioester onto the complement activating pattern (Fig 1A). Activator-bound C3b and factor B (FB) form the alternative pathway (AP) C3 convertase C3bBb ^1,2^ that cleaves C3 and thereby creates an amplification loop (Fig 1A). The AP may initiate independently of pattern recognition by hydrolysis of an internal thioester in C3, converting it to C3(H_2_O) (also known as C3i and iC3) ^3,4^. C3 hydrolysis may also be induced upon contact with biological and artificial surfaces ^5^. The soluble fluid-phase C3 convertase assembles from C3(H_2_O) and FB, where C3(H_2_O)Bb is 2-3 fold less as active than C3bBb ^6–9^. The spontaneous C3 hydrolysis is suggested to maintain a constitutive defense system that opsonizes any surface without host regulators otherwise promoting the dissociation of the convertase C3bBb or degradation of C3b by FI ^10,11^.

**Figure 1.**
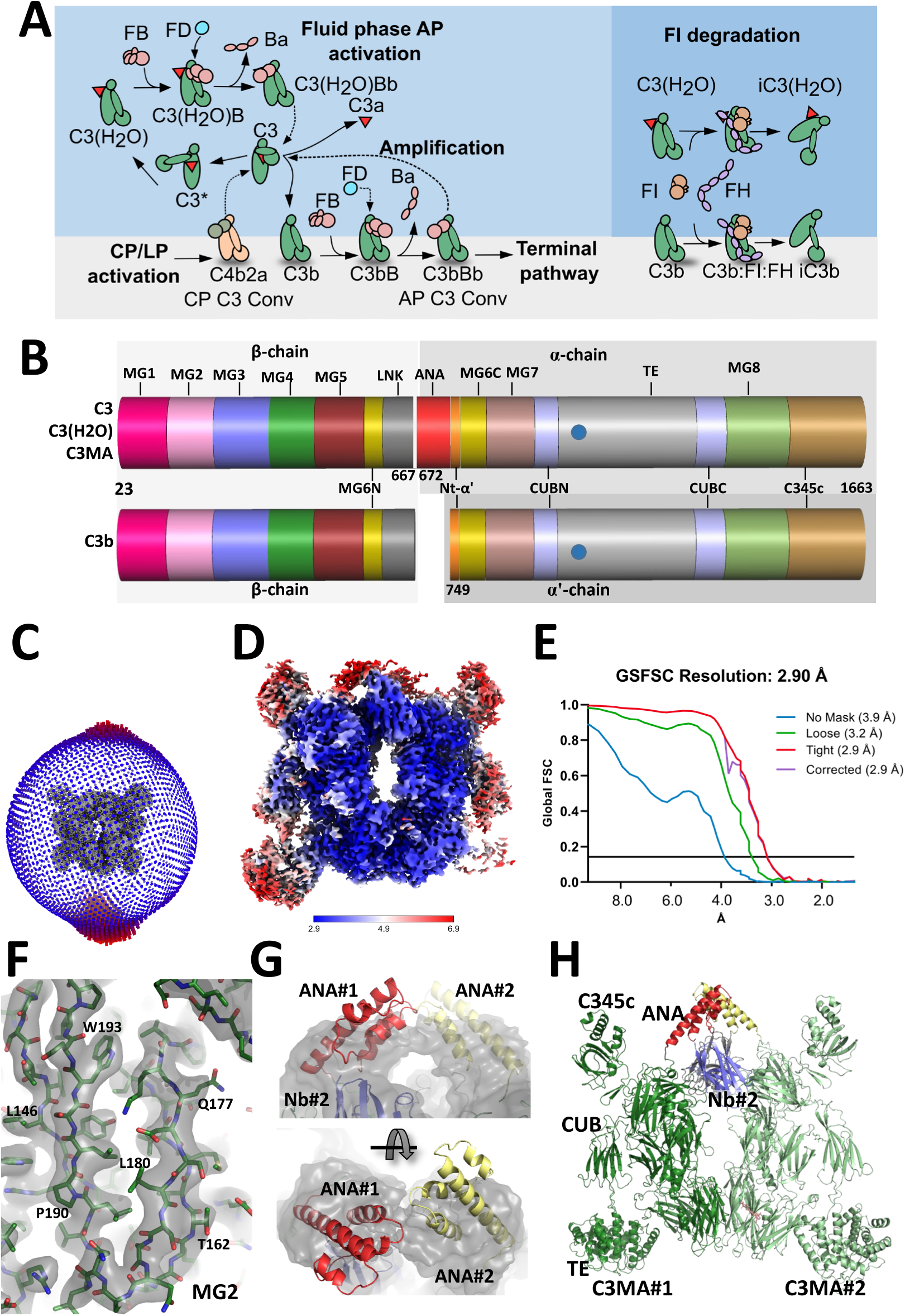
Complement C3 in the alternative pathway and structure determination of C3MA. **A**) C3 is activated by proteolysis to C3b with the release of C3a by either of the C3 convertases C4b2a or C3bBb. C3 is also spontaneously activated by hydrolysis and C3(H_2_O) is generated without C3a release. The fluid phase C3 convertase C3(H_2_O)Bb can generate C3b. In the right part of the figure, the proteolytic inactivation of C3b and C3(H_2_O) by factors I and H is presented. **B**) The chain and domain structure of C3, C3(H_2_O), C3MA and C3b. Note the location of the ANA domain at the N-terminal end of the α-chain. The location of the thioester is marked by a blue circle. **C-D**) The distribution of orientations for the final 3D reconstruction of the C3MA-hC3Nb1 dimer and a local resolution map. **E**) Fourier Shell Correlation (FSC) curve with the FSC=0.143 threshold identified. **F**) Example of the 3D reconstruction for residues in the C3MA MG2 domain contoured at 7σ. **G**) Manual docking of the two ANA domains in a map blurred with a temperature factor of 133 Å^2^ contoured at 3σ. **H**) Structure of the C3MA dimer with the two ANA domains.

Since C3 hydrolysis is slow, methylamine (MA) is used experimentally for aminolysis of the thioester, and MA-treated C3 (C3MA) functionally mimics C3(H_2_O). Both C3MA and C3(H_2_O) form C3 convertase in the presence of complement factors B and D, and both are degraded by factor I in the presence of cofactors like factor H ^9,12,13^. A semi-stable reaction intermediate termed C3* (Fig 1A) with properties distinct from the reactant C3 and the product C3MA/C3(H_2_O) was reported to be structurally unique ^12,14^. The mature C3 protein contains two chains: the N-terminal β-chain and the C-terminal α-chain (Fig 1B). Structures of the 185 kDa protein ^15,16^ have shown that C3 contains the ANA domain, eight macroglobulin (MG) domains, a CUB domain, the thioester containing TE domain, and the C-terminal C345c domain (Fig 1B, see also Supporting Table 1 for definition of the domains). The MG-ring is formed by the MG domains 1-6 and the associated linker (LNK) region. The ANA domain is a helical bundle with 77 residues. Upon proteolytic activation of C3, the scissile bond Arg748-Ser749 between ANA (residues 672-748) and the Nt-α’ region (residues 749-769) is cleaved, and the ANA domain is released as C3a. This triggers a dramatic structural rearrangement in which the TE domain moves by 60 Å ^17^.

Hydrolysis or methylamine treatment converts C3 to C3(H_2_O)/C3MA and does not involve a release of the ANA domain. Yet, C3(H_2_O)/C3MA and C3b are expected to have a similar conformation, as suggested by low resolution structures from negative stain electron microscopy (nsEM) and quantitative cross-linking mass spectrometry (QCLMS) ^14,18^. The MS study concluded that the ANA domain remains on the MG3 domain face of the MG-ring, like in C3. The EM study proposed that the ANA domain passes through the MG-ring to the opposite MG6 domain face ^14^, but the two possible positions of the ANA domain could not be distinguished. Judged by known structures of C3 and C3b ^15–17^, migration of the ANA domain across the MG-ring appears to involve unfolding of the ANA domain or a transient opening of the MG-ring – both requiring significant conformational changes^19^.

## Results

### The ANA domain can migrate through the MG-ring

We set out to determine the structure of C3MA to clarify whether the ANA domain remains on the MG3 face of the MG-ring or migrates to the opposite MG6 face during the C3-C3MA reaction. Here we used the nanobody hC3Nb1 which binds with low nanomolar affinity to C3b, and sterically inhibits progression of the alternative pathway, but does not alter the conformation of C3b ^20^. Although originally found to bind to C3, we recently described that hC3Nb1 does not bind native C3 ^20,21^. Inspired by the crystal packing for a complex between the hC3Nb1 nanobody and C3b ^20^, we mutated Asn108 in the nanobody to cysteine. The resulting disulfide linked nanobody dimer formed highly stable 2:2 complexes with C3MA (Supporting Figure 1A-D). Although this complex formed large single crystals, diffraction was limited, but using cryo-EM single particle analysis (SPA) facilitated by the molecular weight of 400 kDa and approximate two-fold symmetry of the particle, we obtained a 3D reconstruction with a maximum resolution of 2.9 Å of the C3MA-nanobody dimer (Fig 1C-F, Supporting Figures 2 and 3A-B, Supporting Table 2). To verify that the cryo-EM sample contained C3MA and not C3b, we confirmed the presence of ANA by Edman degradation. The retrieved N-terminal sequence corresponded to an intact C3 α-chain starting at position Ser672 (Supporting Figure 1D-E).

**Figure 2.**
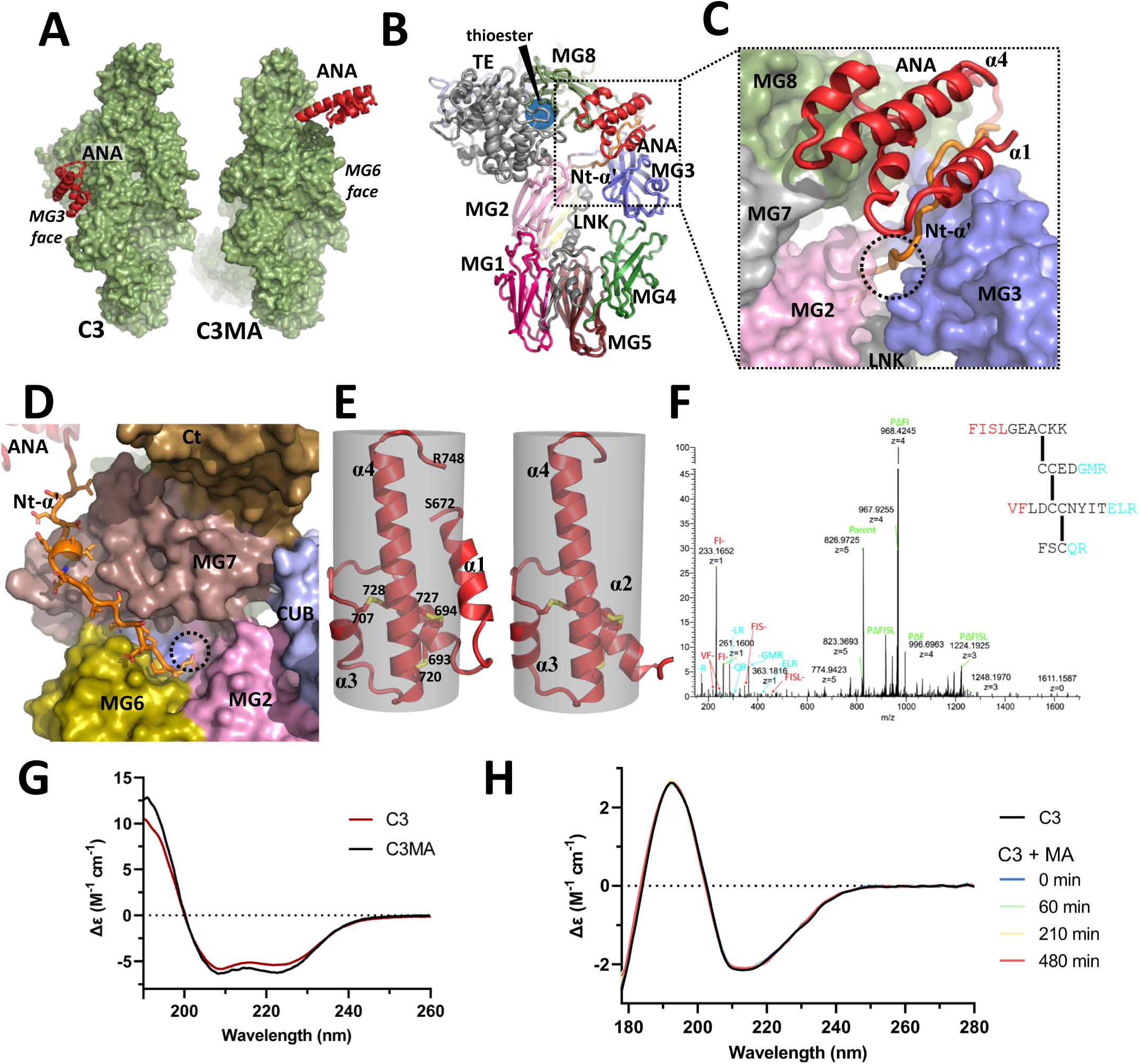
The ANA domain is located at the MG6 face in C3MA, and the secondary structure content is unchanged. **A**) Surface representations of C3 and C3MA in an edge view illustrating how the ANA domain is located on the MG3 face in C3 and on the MG6 face in C3MA. **B-C**) C3 presented from the MG3 face with the domains colored as in Figure 1B. The magnified view illustrates the MG23L channel as a dotted circle (5 Å diameter) between the LNK region and the MG2 and MG3 domains accommodating Nt-α’ residues Ile761-Ser765. **D**) C3MA presented from the MG6 face illustrating how the Nt-α’ is situated in a cleft between the MG6 and MG7 domains, also when the ANA domain is present. The dotted circle indicates the position of the now closed MG23L channel. **E**) Crystal structures of C3a illustrating the four-helix bundle (PDB entry 4hw5, left) and the three-helix bundle (PDB entry 4i6o, right) stabilized by three disulfide bridges enclosed in cylinders of 20 Å diameter. **F**) LC-MS/MS analysis of C3a_T_ resulted in an MS2 spectrum for a tetrapeptide consistent with the known disulfide pattern of C3a, see also Supporting Figure 4B-E and Supporting Table 3. The b-(red) and y-type (blue) product ions and the parent tetrapeptide with and without fragmentation (green) are annotated. **G**) Far UV CD spectra recorded for C3a_T_ released from C3MA and C3 both indicate a high and similar content of α-helices. The minor differences possibly stem from variable methionine oxidation. **H**) SRCD spectra obtained at multiple time points during MA treatment of C3 demonstrate that secondary structure content is maintained throughout the reaction. The experiments in panel F-H were conducted once.

**Figure 3.**
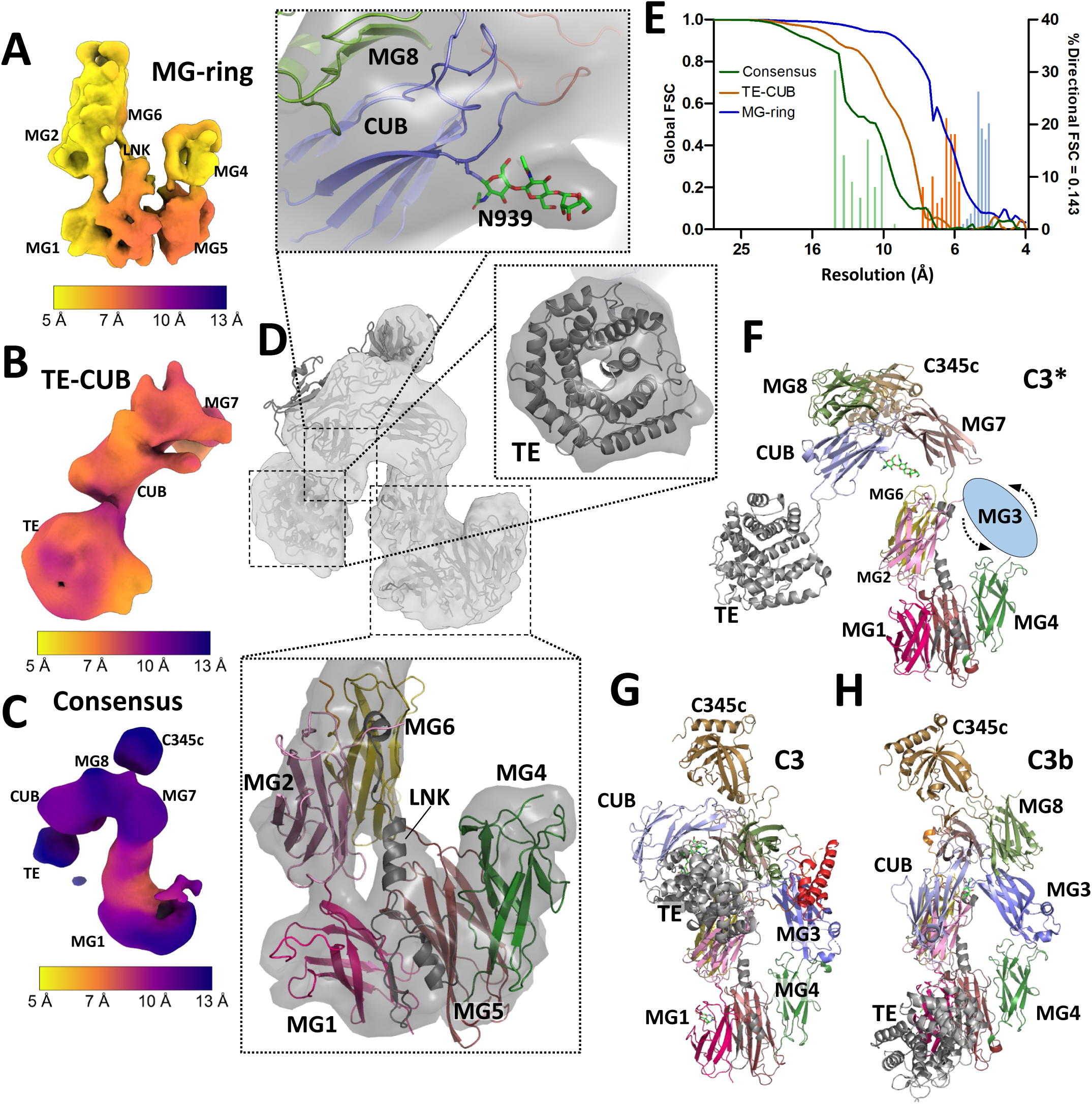
C3* is a highly dynamic molecule with the MG3 domain unlocked from the MG-ring. **A-C)** Three 3D reconstructions of C3* colored by local resolution. The refinement was focused on either the MG-ring (panel A) or the TE and CUB domains (panel B), and a consensus map was obtained by homogeneous refinement of an ab-initio volume centered on MG7 (panel C). **D)** The C3* model superimposed on a map obtained by combination of the three maps in panels A-C. The first three sugar rings of an N-linked glycosylation on Asn939 in the CUB domain can be identified (top), the central hole in the TE domain clearly appears (middle), and a long helix in the LNK region within the MG-ring is recognized (bottom). **E)** Global FSC curves for each of the three maps along with histograms showing the percentage of directional cones with FSC = 0.143 at the given resolution. The global FSCs are determined after mask correction in cryoSPARC, and the directional FSCs have been determined without masking in the 3DFSC program. **F-H**) Comparison of C3*, C3 and C3b. Notice how the TE domain in C3* is located more than 60 Å away from its position in C3 and C3b when structures are aligned on the MG-ring. The model of C3* must be considered hypothetical as it is fitted to the combined map in panel D. In panel F, the blue ellipse marks the dynamic position of the MG3 domain.

From the 3D reconstruction we obtained an atomic model for the entire C3MA except for the ANA domain. Density close to the Nt-α’ region (residues 749-769) that could be attributed to an ANA domain appeared only after blurring of the map, emphasizing the low-resolution features of the 3D reconstruction (Fig 1G). The most likely explanation is that the ANA domain is flexibly attached to the Nt-α’ region, and therefore the 3D reconstruction is noisy at this location. For this reason, the deposited structure of C3MA does not include the ANA domain. Plausible and symmetric locations of the two ANA domains compatible with this blurred density and the connectivity with the ordered Nt-α’ regions were created by manual docking (Fig 1H and Fig 2A).

The overall conformation of C3MA is highly similar to C3b (Supporting Figure 3C-D), consistent with the fact that both can form convertase with FB and be degraded by FI ^22^. The functional similarity between C3MA and C3b was further underscored when we found that C3MA could not compete with iC3b (C3b degradation product) for binding to complement receptor 3 (CR3) (Supporting Figure 3E), in contrast to an earlier report that suggested C3(H_2_O) to be a ligand for CR3 ^23^. In summary, the cryo-EM structure of C3MA showed that ANA together with the Nt-α’ region (residues Ser672-Glu769) is located at the MG6 face of the MG-ring and must have translocated during the conformational change of C3 to C3MA conversion (Fig 2A).

### The structural integrity of the ANA domain is maintained during the C3-C3MA conversion

In C3, the Nt-α’ region threads through a narrow opening delimited by the LNK region together with the MG2 and the MG3 domains (Fig 2B-C). In the following, we refer to this opening as the MG23L channel. In C3MA, the channel is closed (Fig 2D), and the Nt-α’ is located between the MG6 and MG7 domains as seen in structures of C3b and other protease-activated thioester-containing proteins from the α_2_-macroglobulin superfamily ^17,24^. Since the MG23L channel is narrow and only accommodates an extended polypeptide strand in C3 (Fig 2C), it appeared unlikely that the folded ANA domain could pass through this channel during migration across the MG-ring.

We speculated that if the ANA domain unfolds, this may enable its migration across the MG-ring, which also could explain the lack of well-defined density in the cryo-EM map. In C3, the ANA domain is a four-helix bundle stabilized by three disulfide bridges (Fig 2E). To compare the disulfide pattern of the ANA domain in C3 and C3MA, we cleaved both proteins with trypsin. Although C3MA was more susceptible to tryptic cleavage (Supporting Fig 4A), the same major C3a-like fragment corresponding to residues Ser672-Arg740 (called C3a_T_ below since C3a ends at Arg748) was obtained from both C3 and C3MA as determined by LC-MS (Supporting Fig 4B-C, Supporting Table 3). Circular dichroism (CD) spectroscopy analysis of the C3a_T_ from C3 and C3MA revealed two comparable far-UV CD spectra, both with a high α-helical content (Fig 2G). C3a_T_ from C3 and C3MA was further digested by trypsin at pH 6 (to minimize disulfide scrambling) and the resulting peptides were separated by reversed-phase (RP) HPLC, with similar chromatograms obtained for both samples (Supporting Figure 4D-E). The fraction encompassing disulfide-connected peptides was identified using Edman sequencing and further verified by LC-MS/MS. A disulfide-connected tetrapeptide consistent with the established cysteine connectivity was identified in both the C3 and C3MA samples (Fig 2F and Supporting Figure 4D-E). The identification of this tetrapeptide rules out disulfide shuffling except for shuffling between adjacent cysteines. In other experiments, the conversion of C3 to C3MA readily proceeded in the presence of iodoacetamide or mercury ions ^12^, arguing against disulfide shuffling during the reaction. We also observed that C3* progressed to C3MA at pH 6, where disulfide shuffling is minimal (Supporting Figure 4F). In summary, the ANA domain in C3MA had intact disulfide bridges, and if the domain unfolded during migration across the MG-ring, it had subsequently refolded.

**Figure 4.**
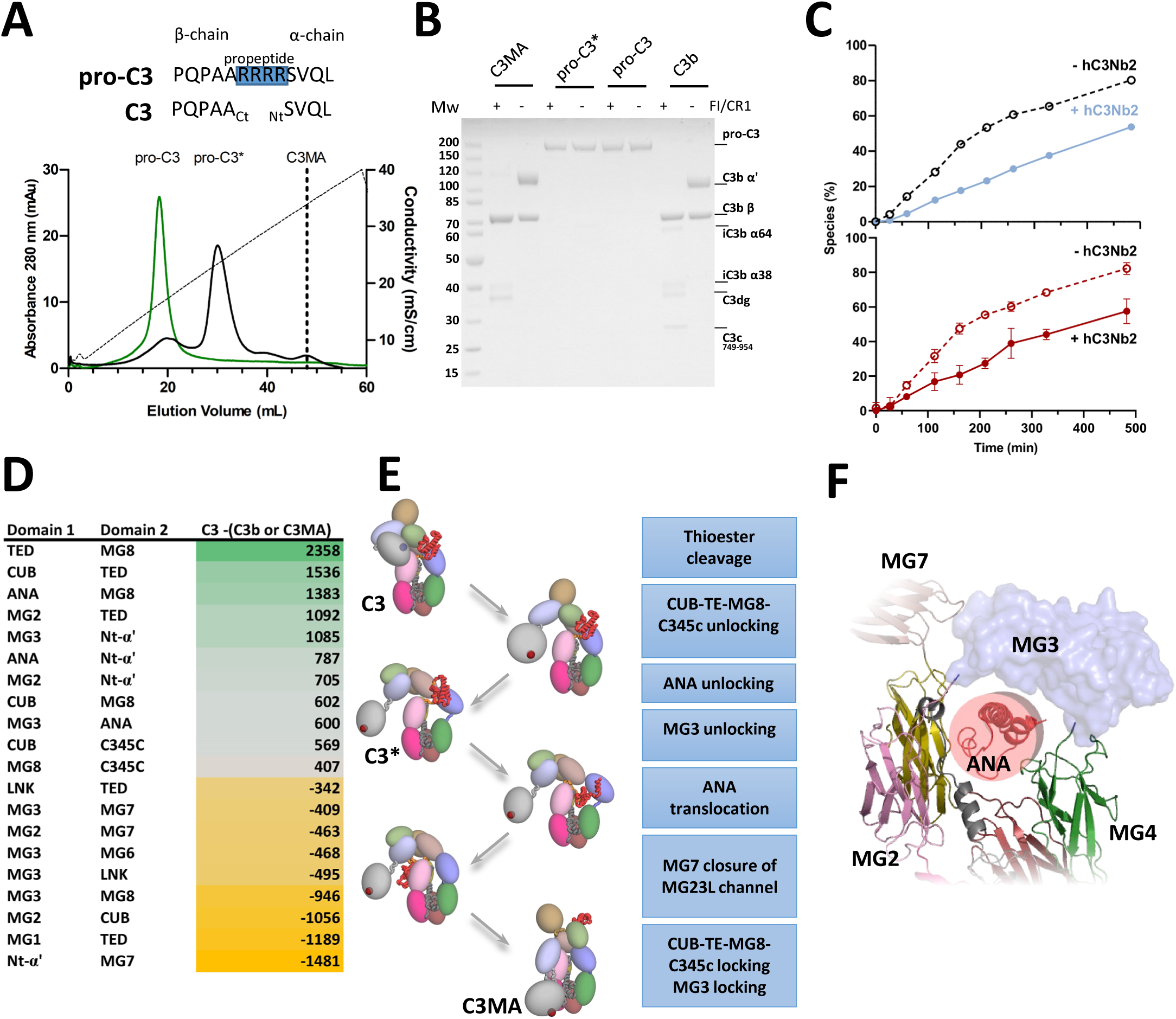
Inhibition and mechanism of the ANA domain migration. **A**) Top: pro-C3 and C3 differ by four residues excised during intracellular maturation. Bottom: Ion exchange analysis of pro-C3 and pro-C3* obtained by 4 hours of methylamine treatment, the elution volume of C3MA is marked for comparison. **B**) Neither pro-C3 nor pro-C3* are degraded by FI, demonstrating that they adopt conformations different from those of C3MA and C3b. **C**) Temporal progression from C3 to C3MA followed by ion exchange (top) or mass-photometry (bottom) with or without the addition of the hC3Nb2 nanobody in three-fold molar excess. Error bars for mass-photometry data show 95% confidence intervals from triplicate measurements. Two biological replications were conducted for the mass-photometry experiment with the parallel ion exchange chromatography measurements performed in one of the two replicates. **D)** A comparison of the buried surface areas of domain interfaces (in Å^2^) in C3 to those in C3b/C3MA. The interaction of the MG3 domain with multiple other MG domains, the interaction of Nt-α’ with MG7 enabled by ANA migration, and the interaction of the TE and CUB domains with the MG-ring appears to drive the conformational change from C3 to C3b/C3MA. **E)** Proposed structure-based mechanisms for nucleophile/hydrolysis induced C3 activation. In the schematic representation to the left, domains are colored as in Figure 1B and the location of the thioester (blue in C3, red in C3*/C3MA) is indicated. The ANA domain, the Nt-α’ and the LNK regions are shown as coils while globular domains are displayed as ellipsoids. **F)** Hypothetical configuration of MG3 and ANA in C3* with ANA in the three-helix bundle conformation. The red cylinder with a diameter of 20 Å illustrates the size of the hole in the MG-ring that may occur in C3* upon MG3 unlocking.

The option remained that ANA or other domains unfold during the C3-C3*-C3MA reaction and refold in the product. A complete loss of secondary structure in the ANA domain is predicted to reduce the content of α-helix from 20% in C3 to 17% in C3* (Supporting Table 1). Unfolding of an MG domain would reduce the content of β-strand from 32% in C3 to 28-29% in C3*. To quantify the content of secondary structure through the reaction, we collected synchrotron radiation CD data from a C3 sample treated with a low concentration of methylamine to induce a slow reaction. The collected spectra were essentially identical, and the calculated α-helix and β-strand content is consistent across C3, C3*, and C3MA (Fig 2H and Supporting Figure 5). The observations from our MS and CD analyses jointly support that neither ANA nor any other domains unfold during the reaction.

### The MG-ring in C3* contains a large dynamic opening

After ruling out significant unfolding of any domains, we used cryo-EM to study the reaction intermediate C3* to reveal changes in the MG-ring during the reaction (Supporting Figure 6 and Supporting Table 4). Single particle analysis of C3* was challenging due to a pronounced conformational variability and preferred orientations, but the MG-ring was recognized in 2D classes from a large data set collected at a tilt angle of 0° (Supporting Figure 6D). To mitigate orientational bias we also included a smaller data set collected with a stage tilt of 25° which vastly improved the 3D reconstructions. Focused refinement with a masked sub volume encompassing the MG-ring resulted in a map where densities for the MG1, MG2, MG4, MG5 and MG6 domains and the LNK region were easily recognized (Fig 3A). Not surprisingly, no density could be attributed to the ANA domain. A rather unexpected finding was that the MG3 domain had no evident density. This implies that in C3*, the position of the MG3 domain relative to the MG-ring is highly variable. Inspection of 2D classes confirmed this notion (Supporting Figure 6D). A second round of local refinement focused on the MG7, CUB and TE domains resulted in a map where these domains were clearly defined with a global resolution of 7.8 Å (Fig 3B and 3E). A third map from unmasked refinement initialized with a volume centered on the MG7 domain, revealed a density that was compatible with the C345c domain linked via the Cys873-Cys1513 disulfide bridge to the MG7 domain and linked to the MG8 domain interacting through a β-sheet with the N-terminal end of the CUB domain (Fig 3C).

A combined focused map derived from these three maps allowed us to construct a model representing C3* (Fig 3D). This map represents a hypothetical state in a spectrum of C3* conformations originating from multiple sources of flexibility: First, the MG3 domain is unlocked from the MG-ring though still tethered between the MG2 and MG4 domains. Second, the MG6-MG7, the MG7-CUB, and to some degree the CUB-TE interfaces are flexible, explaining why no single 3D class presented density for both the MG-ring and the TE domain. Third, the C345c domain appears to have rotational freedom in accordance with its variable location in crystal structures of C3 and C3b ^15–17^.

The model of C3* fitted in the combined map is unique and distinct from any known structures of C3 and C3b (Fig 3F-H). First, the MG-ring is opened due to the unlocking of the MG3 domain. Second, the TE domain is completely exposed at the end of the CUB domain and located more than 60 Å from the positions in C3 and C3b (Fig 3F-H). In the C3* model (Fig 3F), the MG8 domain is apparently located more than 40 Å from its position in C3 and C3b but the quality of the density suggests a variable position. The MG7 domain is located above the MG2 and MG6 domains in a unique location compared to both C3 and C3b (Fig 3F).

Since the C3* conformation was derived from a map assembled from multiple low-resolution 3D reconstructions, we also studied the conformation of C3* in solution using small angle X-ray scattering (SAXS) at pH 6.0. Without any model optimization, our C3* cryo-EM model both qualitatively explained the distance distribution function and fitted the experimental data much better than models of C3 and C3MA (Supporting Figure 7). Placing the MG3 and ANA domains oriented as in native C3 into the C3* model from cryo-EM deteriorated the fit, thus providing further evidence of a dynamic association of the MG3 domain with the MG-ring. Upon rigid body optimization where the MG3 and ANA domains were allowed to move relative to the rest of the C3* model, a model was produced with an improved fit (χ^2^ = 1.4) of the calculated scattering curve to the observed data (Supporting Figure 7).

In summary, the observed flexibility of C3* and the unexpected unlocking of the MG3 domain provided evidence that the MG-ring temporarily opens in C3*. This could provide a route for migration of the ANA domain across the MG-ring.

### ANA migration and MG3 unlocking are required for formation of C3MA

Pro-C3 is a single chain peptide, which furin cleaves to generate the C3 α- and β-chains prior to secretion (Fig 4A). Since the ANA domain is at the N-terminal end of the α-chain at the MG3 face, in pro-C3 the ANA domain should be unable to migrate across the MG-ring to the MG6 face. To confirm this idea, we prepared a recombinant form of pro-C3, in which the furin cleavage site (residues 668-671) was mutated to prevent maturation. In nsEM, this pro-C3 resembled native C3 (Supporting Figure 8A). Upon treatment with MA, pro-C3* appeared, demonstrating that in pro-C3 the thioester is formed and protected as in C3 (Fig 4A). As judged from our chromatographic analysis, almost no pro-C3MA was formed after 4h (Fig 4A) indicating that the ANA migration could not take place or was much slower in our recombinant pro-C3. In addition, neither pro-C3 nor pro-C3* could be cleaved by FI, further supporting that they do not adopt a C3b/C3MA-like conformation (Fig 4B). To our knowledge, this is the first example of a C3 variant that cannot reach the activated conformation of C3MA but instead accumulates in the C3* conformation.

In the observed C3* conformation, the TE domain is highly exposed compared to known structures of C3 and C3b/C3MA (Fig 3F-H). To investigate whether the TE domain is required for formation of the C3* conformation, we prepared the variant C3ΔTE in which the TE domain is deleted. To monitor its conformation, we conducted an FI degradation assay with a CR1 fragment as a cofactor and observed degradation of the C3ΔTE α-chain missing the TE domain (Supporting Figure 9A-B). This suggested that in C3ΔTE, the CUB domain may be located next to the MG2 domain, and that the MG-ring adopts a C3b/C3MA like conformation. This was confirmed by negative stain EM analysis of C3ΔTE (Supporting Figure 9C-E).

Overall, the experiments with C3ΔTE suggested that the C3* conformation depends on the presence of the TE domain and that this semi-stable conformational state most likely only can be reached through nucleophilic activation starting from the conformation of native C3 with an intact thioester. To our knowledge, our data on C3ΔTE demonstrates for the first time that the C3b/C3MA conformation of the MG-ring and its interaction with the CUB domain may occur independently of the TE domain.

If the MG3 domain is unlocked in the C3* state as suggested by cryo-EM, an MG3 binding molecule may influence the formation of C3MA. We therefore measured the rate of C3MA formation in the presence and absence of the MG3/MG4-interface-binding nanobody, hC3Nb2 ^25^, and followed the transformation using ion exchange chromatography. In an orthogonal approach, we took advantage of the tight interaction between C3MA and the EWEµH fusion protein ^21^ to follow the formation of the C3MA by mass photometry in the absence or presence of hC3Nb2. The EWEµH encompasses a modified version of hC3Nb1 called EWE fused with a linker to the FH fragment CCP2-4. The EWE moiety binds at the same place as hC3Nb1 while the FH fragment is likely to bind as known from the C3b-FI-FH complex ^26^. Both methods consistently demonstrated that a three-fold molar excess of hC3Nb2 to C3 decreased the rate of C3MA formation roughly two-fold in the early phase of the reaction (Fig 4C and Supporting Figure 10). Overall, the data obtained with the hC3Nb2 nanobody provided compelling evidence that the MG3 domain has a key role in the conversion of C3 to C3MA as suggested by our C3* structure.

Since we observed that the hC3Nb2 nanobody inhibited C3MA formation, we investigated C3* in complex with hC3Nb2 by cryo-EM following the same experimental strategy used for the C3*. We observed a closed MG-ring in which the MG3 domain with bound hC3Nb2 became visible in both 2D classes and a 3D reconstruction at 5.1 Å resolution (Supporting Figure 11 and Supporting Table 5). Hence, the nanobody stabilizes the position of the MG3 domain within the MG-ring in the C3* conformation, and this explains the observed inhibition of C3MA formation by hC3Nb2 (Supporting Figure 10).

We also evaluated the rate of C3MA formation under influence of monomeric hC3Nb1 nanobody, using our cation exchange chromatography assay for quantitation of C3, C3* and C3MA. In contrast to the inhibition observed for hC3Nb2, we observed a significant stimulation of the rate of C3MA formation and specifically much less accumulation of C3* as compared to the unstimulated reaction (Supporting Figure 12). In the absence of hC3Nb1, the amount of C3* was higher or equal to that of C3 at all time points except for t = 0 h, whereas in the presence of hC3Nb1, C3* was lower or equal to the amount of C3 at all time points (Supporting Figure 12B). This may be explained if a late intermediate in the C3* to C3MA reaction binds hC3Nb1 and that this interaction promotes the irreversible formation of the C3MA product.

### Mechanism for C3 activation by nucleophiles

Both proteolytic and nucleophilic activation of C3 results in highly similar conformations of the respective products C3b and C3MA (Supporting Figure 3C-D). A comparison of the buried surface areas in pairs of domain interactions in C3 and C3b/C3MA emphasized the high similarity (Fig 4D and Supporting Table 5). Proteolytic activation to C3b is fast and the ANA domain is released whereas nucleophilic activation through thioester breakage is slow ^22^ and the ANA domain is retained and translocated through the MG-ring. Proteolytic activation occurs by cleavage of the scissile bond Arg748-Ser749 located 50 Å from the thioester where nucleophile activation starts (Fig 2B). Therefore, the order and timing of smaller steps in the conformational changes occurring in the two systems must be different.

The first steps in the nucleophile-induced, slow conformational change must be thioester cleavage and disruption of the MG8-TE domain interface (buried surface area of 2358 Å^2^) that is key to the conformation of native C3 and the protection of the thioester from hydrolysis ^15,16^. In C3, the ANA domain is located between the MG3 and the MG8 domains (Fig 2C), consequently, the release of the MG8 domain from its interaction with the TE domain, is likely to lead to disruption of the MG8-ANA domain interface (1383 Å^2^). We suggest that this is followed by unlocking of the ANA and MG3 domains (Fig 4E). The result is the long-lived dynamic intermediate observed in our C3* structure. The irreversible conversion of this intermediate to the product can only occur after ANA migrates to the MG6-face of the MG-ring. The process of ANA migration through the MG-ring enabled by the MG3 unlocking is a very plausible candidate for the rate-limiting step of the reaction and maybe the step stimulated by hC3Nb1. We suggest that the subsequent movement of the MG7 domain is driven primarily by its interaction with the now relocated Nt-α’ region (1481 Å^2^). Following the MG7 movement, the conformational change that places the TE domain next to the MG1 domain and the CUB domain next to the MG2 domain can reach completion and the tight interactions between the MG3 domain and the MG8 (1481 Å^2^) and the MG7 domains (806 Å^2^) that firmly lock the MG-ring in the well-known C3b conformation can be established. This proposed flow of events for the MA activation of C3 is in accordance with a prior three-step model based on spectroscopy ^22^.

The migration of ANA through the MG-ring is an unusual and necessarily slow molecular event. In C3*, the extended residues Nt-α’ (residues 749-769) connecting the ANA domain with the MG6 domain may easily pass through the MG23L channel akin to the translocation of the Nt-α’ region in nascent C3b, but a folded ANA must pass the MG-ring through a wider opening. When the MG3 domain is dynamically suspended between the MG2 and the MG4 domains as directly captured in our structure of C3*, a much larger opening than the MG23L channel becomes possible but is still unlikely to be significantly wider than the smallest dimension of the ANA domain (Fig 4F). A three helix bundle conformation of C3a with the same disulfide bridge pattern as the four helix bundle is known ^27^ . The ANA domain is likely to adopt this three-helix bundle conformation in pro-C3 as the four residues in the propeptide are unable to span the distance between Ala667 and Ser672 in the four-helix bundle conformation of ANA known from C3 (Supporting Figure 8B). An ANA domain in the three-helix bundle conformation may pass more easily through the hole in the MG-ring that we observe in C3* compared to a four-helix bundle (Fig 4F). We also note that in native C3, one MG3 β-sheet interacts with residues in the Nt-α’ and the first α-helix in the ANA domain. If such interactions also transiently occur in C3*, they could increase the probability of the ANA domain slipping through the MG-ring as compared to the case where the ANA and MG3 domains move independently by Brownian motion. Two short helices in C3 (residues Lys600-Val620) located in the LNK region at the MG3 face of the MG-ring may also play a role during the migration of ANA in accordance with a higher mobility of this region in C3 compared to C3b suggested by hydrogen-deuterium exchange ^19^.

In summary, we propose that a nucleophile attack on the thioester in C3 causes the release of the MG3 domain from the MG-ring, possibly together with local conformational changes of the LNK region. In this way, the intermediate C3* has an opening large enough for the folded ANA domain to slip through in a slow, rate-limiting step (Fig 4F). Once ANA has migrated to the opposite face of the MG-ring, movement of the MG7 domain allows the CUB, TE, MG8 and C345c domains to settle to the positions also known from C3b. The re-integration of the MG3 domain into the MG-ring is likely to be secured by substantial interactions with the MG8 domain as observed in both C3MA and C3b (Supporting Figure 3D).

## Discussion

Our cryo-EM structure of C3MA directly shows that the ANA domain is located at the MG6 face of the MG-ring and therefore must be able to pass from the MG3 face where it resides in native C3. This was expected since C3MA, like C3b, can form convertase and is degraded by FI. However, upon comparison of C3MA/C3b and C3 structures, it was difficult to understand the mechanism for ANA translocation due to the limited aperture of the MG23L channel, i.e. there was no obvious path for the migration of ANA without major structural rearrangements. Unfolding of the MG6 domain was previously proposed to allow migration of the ANA domain across the MG-ring ^14^ ^19^, but our data strongly implicates the MG3 domain as the key player in the MG-ring. Our SRCD and MS data imply that ANA migration occurs without major loss of secondary structure or disulfide rearrangement. Furthermore, if significant unfolding occurred in the MG-ring of C3*, it would have to be reversible as the structures of the MG-ring in C3MA and C3b are indistinguishable (Supporting Figure 3C-D). One limitation is that CD spectroscopy is unable to register temporary loss of secondary structure in a minor fraction of molecules. Therefore, we cannot exclude that translocation of the ANA domain across the MG-ring involves transient breakdown of secondary structure or collapse of tertiary structure in the slow, rate-limiting step C3*-C3MA. However, the unexpected opening of the MG-ring created by the mobility of the MG3 domain in C3* argues that dramatic changes in the secondary structure of ANA and MG domains are not required. It remains an option, however, that the ANA and the MG3 domains to some degree reversibly change their tertiary structure or enter a molten globule-like state ^28^ in the rate-limiting step.

A prior QCLMS study suggested a C3b-like structure of C3(H_2_O), where the ANA domain was found to cross-link with the MG3, MG7 and MG8 domains, and it was concluded that ANA does not migrate through the MG-ring ^18^. However, the proximity of ANA to these MG domains is compatible with our cryo-EM structure of C3MA considering that the hC3Nb1 nanobody keeps ANA suspended above the MG7 and MG8 domains. Conformational dynamics of the extended Nt-α’ region may enable the ANA cross-links observed by QCLMS.

The conformational change taking place in the C3-C3MA reaction is exceptional compared to that occurring during the C3-C3b reaction ^17^. As one measure, the center of mass of the TE domain moves by 60 Å in the C3-C3b transition. In C3*, the CUB-TE domain tandem is rocking with the MG7 domain and in the combined map representing a single hypothetical C3* conformation (Fig 3D and 3F), the TE domain is roughly 60 Å from its distinct locations in C3 and C3b. The conformational change occurring in the C3-C3MA transition is also unique in size and dynamic nature compared to other macromolecules. This is illustrated by comparison with serpins. These protease inhibitors are, like C3, cleaved by a protease and undergo a fast, large-scale conformational change involving insertion of the cleaved reactive center loop in a β-sheet, formation of a covalent acyl-enzyme intermediate, and disruption of the protease active site geometry ^29^. Furthermore, in parallel to the high structural similarity between C3b generated by proteolysis and C3MA obtained by thioester cleavage, some serpins and, in particular, plasminogen activator inhibitor type 1 (PAI-1) spontaneously rearrange to the latent form that shares the insertion of the reactive center loop in a β-sheet with the protease activated serpins ^30^. Compared to this well-established active-to-latent transition in serpins, the irreversible conformational change induced by C3 thioester breakdown that we report here is unique in scale, dynamic nature of the C3* intermediate, and mechanism.

In humans, C3 is present at 6.5 μM (1.2 mg/ml) in plasma ^31,32^ and in a 70 kg individual, the daily turnover of C3 is more than 200 mg at a putative tick-over rate of 0.3% per hour ^3^. Accordingly, in fresh serum and plasma samples, the concentration of C3(H_2_O) is ∼500 ng/ml ^33^. C3 hydrolysis is widely presented as a third initiation pathway for C3b deposition without prior activation of the classical and lectin pathway ^34,35^, but the ability of C3(H_2_O) to form an efficient fluid phase convertase in healthy individuals has been challenged ^10,11^. In the fluid phase, FI assisted by FH rapidly degrades C3(H_2_O) to iC3(H_2_O), which is unable to form convertase (Fig 1A) ^9,36^. Fluid phase C3b generated by C3(H_2_O)Bb may also be inactivated prior to reaching a cell where it can attach covalently ^4^. Nevertheless, C3(H_2_O) could still be an important functional state, as C3(H_2_O) and iC3(H_2_O) were suggested to act as sources of intracellular C3a. Multiple cell types including B- and T-cells take up extracellular C3(H_2_O)/iC3(H_2_O) and recycle it to the extracellular environment, and in CD4^+^ T cells this recycling influences the cytokine profile ^37^. C3(H_2_O)/iC3(H_2_O) was also found on platelets and shown to be involved in platelet-leukocyte interactions ^23^.

A higher rate of hydrolysis is induced by C3 contact with biological and artificial surfaces ^5^ with C4-independent C3 activation during hemodialysis ^38^ and C3 activation at the gas-plasma interface in relation to air embolism ^39^ as prominent examples. Recent studies also link C3(H_2_O) to heme-mediated complement activation that induces injury to endothelial cells, *in vitro* and *in vivo* models of intravascular hemolysis ^40,41^. *In vivo*, FH assisted FI regulation of C3(H_2_O) is crucial as demonstrated by the variant C3-923_ΔDG_ that causes dense deposit disease ^42^. Recently, high levels of S-nitrosylated cysteine residues in C3 were detected in brains from especially females with Alzheimer’s disease, and this modified C3 was linked to increased phagocytosis, synapse loss and cognitive decline ^43^. In particular, Cys720 in the ANA domain was S-nitrosylated, and we speculate that this destabilizes the tertiary structure of the ANA domain and alters the rate of C3 hydrolysis and the properties of C3*.

Numerous therapeutic strategies targeting the alternative pathway have been developed to treat diseases like age-related macular degeneration, paroxysmal nocturnal hemoglobinuria, atypical hemolytic syndrome, and C3 glomerulopathy ^44,45^. Multiple C3 inhibitors are known including the compstatin like inhibitors ^46,47^, but none of these strategies selectively target C3(H_2_O), and the lack of selective and cross-reactive C3(H_2_O) inhibitors prevents *in vivo* evaluation of the contribution of C3 hydrolysis to complement activation. Our experiments with pro-C3 and hC3Nb2, where the C3-C3MA transition is either prevented or delayed, outline for the first time two different approaches to interfere with C3 hydrolysis. Our structures and functional studies of C3* and C3MA offer a rational framework for the development of highly specific inhibitors of C3(H_2_O). Considering the prompt inactivation of C3(H_2_O), C3* could be significantly more abundant than C3(H_2_O) and therefore be a relevant target for the development of a complement inhibitor.

## Materials and methods

### Preparation of proteins

Human native C3 and C3b were prepared as described ^48^. C3MA and C3* for cryo-EM were generated from purified native C3 in 20 mM HEPES pH 7.5, 200 mM NaCl. The reaction was performed at 200 mM CH_3_NH_2_ (methylamine, MA) pH 8.5 and 10 mM iodoacetamide, followed by incubation at 37 °C on a tabletop shaker. To obtain C3*, the reaction ran for 15 minutes. To obtain C3MA, the reaction ran for at least 4 h. After incubation, the sample was moved to ice, pH adjusted with 1 M MES pH 6.0 and diluted to 60 mM NaCl. Next, the sample was loaded onto a 1 mL Mono S column (Cytiva) equilibrated in 20 mM MES pH 6.0, 50 mM NaCl. The C3 species were separated with a 60 mL linear gradient from 50-400 mM NaCl. Biotinylated iC3b and the complement receptor 3 headpiece (CR3HP) were generated as described ^49^. Pro-C3 was expressed as described ^50^ from a plasmid where the arginine residues 668-RRRR-671 were mutated to the sequence GSGR. Secreted pro-C3 was purified from the HEK293 cell culture supernatant. After filtration through a 0.22 µm filter, the supernatant was supplemented with Tris-HCl pH 7.5 to 50 mM and diluted three-fold in 50 mM Tris-HCl pH 7.5. Finally, 0.5 mM PMSF and 5 mM EDTA were added. The supernatant was loaded on a 5 mL HiTrap Q FF column (Cytiva) and the column was washed with 20 mM Tris pH 7.5, 50 mM NaCl. The pro-C3 was eluted with a 30 mL linear gradient from 50-500 mM NaCl. Fractions containing pro-C3 were pooled and dialyzed against 20 mM MES, 100 mM NaCl, pH 6 at 4 °C. Before loading on a 1 mL Mono S column equilibrated in 20 mM MES pH 6.0, 50 mM NaCl, the sample was diluted with 20 mM MES pH 6 to 65 mM NaCl. The protein was eluted by a 30 mL linear gradient from 50-400 mM NaCl. Peak fractions were analyzed on SDS-PAGE. The hC3Nb2 nanobody and the EWEµH fusion proteins were prepared as described ^21,25^. C3ΔTE was expressed from a modified C3 expression plasmid where the residues 990-1287 were deleted and residues 989 and 1286 connected. This C3 variant was expressed and purified as pro-C3 but was purified on a 1 mL Source 15S column instead of a 1 mL Mono S column.

### Preparation of the hC3Nb1 dimer

To introduce the N108C mutation in the expression plasmid for the parental hC3Nb1 ^20^, mutagenesis was performed using the QuickChange® Lightning Site-Directed Mutagenesis Kit (Agilent) using the forward primer 5’ CAAACAGATTGTGAGTATAACTACTGGGGCCAG 3’ and the reverse primer 5’ AGTTATACTCACAATCTGTTTGTGGACTCCAACCCG 3’. *E. coli* cells expressing hC3Nb1 N108C were grown to OD600 ∼ 0.8 in 2×TY medium and cooled on ice followed by a 30-minute incubation at 20 °C and expression induced by addition of 1 mM IPTG. After overnight expression at 20 °C, the cells were harvested and hC3Nb1 N108C was purified as described ^20^. For dimerization, purified hC3Nb1 N108C was added a 10-fold molar excess of DTNB (Sigma-Aldrich) and allowed to react for 30 minutes at room temperature, followed by buffer exchange into 20 mM HEPES pH 7.5, 150 mM NaCl on a PD10 column (Cytiva). Next 0.95 equivalents of unreacted hC3Nb1 N108C was added to the monomeric hC3Nb1 N108C DTNB adduct for formation of the dimer. The reaction was left at room temperature for 10 h and then diluted fivefold in 20 mM sodium acetate pH 5.5. The sample was loaded on a 1 mL Source 15S column (Cytiva) equilibrated in 20 mM sodium acetate pH 5.5, 150 mM NaCl. hC3Nb1 N108C dimer and monomer were separated by a 40 mL linear gradient from 150-450 mM NaCl.

### Large scale trypsin digestion of C3MA and C3 for CD and MS analysis of C3a_T_

Trypsin digestion of C3 or C3MA was performed in 1 ml reactions with protein present at 2 mg/ml in 20 mM HEPES pH 7.5, 100 mM NaCl with 10 mM iodoacetamide (Sigma-Aldrich). The digestion was started by addition of 4 µl 0.1 mg/ml trypsin (Sigma-Aldrich T8003) to each 1 ml reaction. The reaction mixture was incubated at 37 °C for 6 min on a tabletop shaker. The reaction was stopped by moving to ice and adding the protease inhibitor Pefabloc (Roche) to a final concentration of 1 mM. Freshly trypsinated C3 or C3MA was diluted 1:1 with 20 mM MES pH 6.0 and next 50 mM MES pH 6.0 was added from a 1 M stock solution. After filtration, the sample was loaded on a 9 ml Source 15S (Cytiva) column equilibrated in 20 mM MES pH 6, 30 mM NaCl. The products C3b and C3a_T_ were eluted with a gradient from 30-350 mM NaCl over 70 ml. The C3a_T_ containing fractions were pooled and adjusted to final 20 mM HEPES pH 7.5. Before CD measurement, a final SEC purification step and buffer exchange was performed on a 24 ml Superdex 75 increase column (Cytiva) equilibrated in 137 mM NaF, 10mM Na_2_HPO_4_ pH 7.4.

### MS analysis

To study the mass of C3a_T_ prepared from native C3 and C3MA, the C3a_T_ samples were desalted using pipette tips packed with POROS 50 R2 C18 resin (PerSeptive Biosystems). Then, ∼25 nanograms of C3a_T_ were analyzed by LC-MS/MS with an EASY-nLC 1200 (Thermo Fisher Scientific) and an Orbitrap Eclipse Tribrid mass spectrometer (Thermo Fisher Scientific). A data-dependent acquisition method selected peptides for fragmentation by high-energy collision dissociation (HCD) and MS2; only MS1 spectrum data were used for intact mass determination. Data analysis was performed with the FreeStyle software (Thermo Scientific, version 1.8.63.0), which was used to deconvolute an average MS1 spectrum that included all eluting peptides. The deconvoluted masses were manually compared to the C3a sequence in residues 672-748 of UNIPROT entry CO3_HUMAN using the GPMAW software (Lighthouse Data, version 9.51) to identify cleavage sites for trypsin in the generation of C3a. The oxidation of methionine and the formation of disulfides from cysteines were the only amino acid modifications considered in the analysis.

To compare the disulfide pattern of C3a_T_ prepared from native C3 and C3MA, the C3a_T_ samples were digested with a 1:10 w/w ratio of trypsin:C3a_T_ using MS-grade trypsin overnight at 37 °C in 50 mM MES pH 6 (to minimize disulfide scrambling). The tryptic peptides were then separated by offline RP-HPLC using a Brownlee Aquapore RP-300 column and a gradient from 0-90% acetonitrile, with 0.1% TFA. The sequences of peptides in the highest-intensity peaks were analyzed by Edman sequencing, and a peak with a disulfide-containing peptide from C3a_T_ was identified in each sample. The fraction containing this peak was prepared for MS analysis by desalting using pipette tips packed with POROS 50 R2 C18 resin (PerSeptive Biosystems). Then, approximately 100 nanograms of peptide were analyzed by LC-MS/MS with an EASY-nLC 1200 (Thermo Fisher Scientific) and an Orbitrap Eclipse Tribrid mass spectrometer (Thermo Fisher Scientific) running a data-dependent acquisition method, where selected peptides were fragmented for MS2 by high-energy collision dissociation (HCD). Assignment and annotation of MS2 spectra were performed manually, considering only disulfide formation as an amino acid modification.

### CD data collection and analysis

Far-UV circular dichroism spectra on C3a_T_ from C3MA were recorded on a Jasco J-810 spectropolarimeter at 293 K in a 1 mm path-length quartz cuvette. The spectrum was recorded with a scan speed of 20 nm/min and a response time of 2 s over 5 accumulations. Spectra recorded of C3a_T_ from native C3 were recorded on a Jasco J-815 spectropolarimeter, also at 293 K in a 1 mm path-length quartz cuvette: however, with a scan speed of 50 nm/min and a response time of 1 s over 40 accumulations. In both cases C3a_T_ was present at a concentration of 12 µM in a buffer containing 10 mM NaH_2_PO_4_ pH 7.4, 137 mM NaF. A conclusive concentration determination on a UV-VIS spectrophotometer was performed immediately after acquiring CD-spectra. A blank buffer spectrum used for background subtraction was recorded for each sample with identical parameters.

To enable CD analysis of the MA reaction, purified C3 was dialyzed overnight into 30 mM Na_2_HPO_4_ pH 8.0 and diluted to 0.8 mg/mL. To initiate the reaction, 10 mM methylamine pH 8.0 was added to the mixture and incubated at 37 °C on a shaker operated at 180 rpm. Samples were removed at different time-points (0, 25, 60, 115, 160, 210, 260, 330 and 480 min) and used for Synchrotron Radiation Circular Dichroism (SRCD) spectroscopy and for cationic exchange chromatography analysis. At each time point, 20 µL of the reaction was loaded into a 0.01 cm path length quartz cell (SUPRASIL, Hellma GmbH, Germany) for SRCD measurements. Each spectrum was recorded on the AU-CD beam line at the ASTRID2 synchrotron radiation source (ISA, Aarhus University, Denmark) with 1 nm steps and a dwell time of 2.1 s per step between 170-280 nm. Each time point was measured 6 times, and since no significant differences were observed the spectra were averaged out. Δε was calculated using protein concentration estimated from the absorbance at 205 nm. The secondary structure content was evaluated using DichroWeb analysis tools ^51^ with CDSSTR as the analysis program and using the SP175 dataset as reference. The results of the best ten predictions for the secondary structure were averaged, and the 95% confidence interval from these measurements was calculated.

In parallel, the reaction was followed by cationic exchange chromatography. For each time point, 50 µg of MA-treated C3 was diluted 20-fold in 10 mM MES pH 6.0 and loaded on a 1 mL Source 15S column equilibrated with 10 mM MES pH 6.0, 50 mM NaCl and eluted with a gradient from 50 to 300 mM NaCl over 15 mL. Pure C3, C3* and C3MA were used as controls to identify the elution volumes of the three molecules. The fraction of each species during the reaction was deconvoluted with PeakFit (SYSTAT) using the elution volumes obtained from the controls.

### C3-C3MA time course experiments

To follow the generation of C3* and C3MA at pH 8.5, 1 mg of purified C3 in 20 mM HEPES pH 7.5, 150 mM NaCl was treated with 200 mM methylamine, 20 mM Tris pH 8.5, and 10 mM iodoacetamide. The reaction was incubated at 37 °C in a shaker at 450 rpm. Samples were taken at 0, 30, 90, 180, and 240 min, and the reaction was stopped by moving the sample to ice and adjusting to pH 6 by addition of 1 M MES pH 6.0 to a final concentration of 50 mM. The sample was further diluted in 20 mM MES pH 6.0 to reach a concentration of 60 mM NaCl. Next, the sample was loaded onto a 1 mL Mono S column equilibrated in 20 mM MES pH 6.0, 50 mM NaCl. The C3 species were eluted by a 60 mL linear gradient from 50-400 mM NaCl. To examine the ability of C3* to form C3MA at pH 6.0, purified C3* was diluted 1:3 with 10 mM MES pH 6.0 and incubated for 30 min at 37 °C with shaking. After incubation, the sample was analyzed by cation exchange chromatography on a 1 ml Source 15S column using a gradient from 50-300 mM NaCl in 10 mM MES pH 6.0.

### C3MA formation in the presence of the hC3Nb2 nanobody

Native C3 was dialyzed into 30 mM Na_2_HPO_4_ pH 8.0 and diluted to 0.8 mg/mL, either with or without a 3-fold excess of hC3Nb2. After addition of 10 mM methylamine, the sample was incubated at 37 °C in a shaker. Samples were taken at time points 0, 25, 60, 115, 160, 210, 260, 330 and 480 min and transferred to ice. For mass-photometry analysis on a TwoMP instrument (Refeyn), 2 μL sample from each time point was diluted to 1 μM total C3 and either mixed with a two-fold molar excess of EWEμH and incubated on ice for 5 min or measured directly after dilution to a final 5 nM C3. Samples mixed with EWEμH were diluted to approximately 3 nM of C3. Each mass-photometry measurement was performed for 1 min in triplicates. Calibration measurements of ovalbumin, aldolase, ferritin and alpha-2-macroglobulin were done at the beginning, after 4 hours and after 8 hours, and their combined calibration curve was used for all samples.

To estimate the abundance of C3MA in the sample, the fraction of EWEμH-bound events compared to total C3 events was estimated by Gaussian fitting both peaks in the Refeyn DiscoverMP software when peaks were properly resolved. If the peaks were not resolved, C3MA abundance was determined from counting events in the shoulder. This was done by comparison to a control measurement without addition of EWEμH by normalizing the control peak to the peak height of the sample measurement. Bins next to the peak with consecutively higher counts in the sample than in the control were summed and counted as the shoulder. The estimate of C3MA-abundance was obtained by normalizing the fraction of EWEμH-bound events to those found in control measurements with C3MA.

In parallel, and similarly to the SRCD experiment (now in the presence of hC3Nb2), 50 µg of reacted C3 was diluted 20-fold in 10 mM MES pH 6.0 and loaded into a Source 15S 1 mL pre-equilibrated with 10 mM MES pH 6.0, 50 mM NaCl buffer and eluted with a linear gradient from 50 to 375 mM NaCl over 20 mL. The interaction between hC3Nb2 and C3 species increased the conductivity at which the complexes eluted.

Therefore, control samples of C3, C3* and C3MA were incubated with a 3-fold excess of hC3Nb2 to identify their elution volumes. This showed that C3 and C3* complexed with hC3Nb2 elute closer to each other in comparison with free C3 and C3*, and therefore difficult to distinguish and deconvolute. The elution volumes obtained from the controls were used for PeakFit deconvolution of hC3Nb2 and C3MA:hC3Nb2 from C3:hC3Nb2/C3*:hC3Nb2. The elution volume and the full width at half maximum (FWHM) was kept fixed for hC3Nb2 and C3MA:hC3Nb2.

### C3MA formation in the presence of the hC3Nb1 nanobody

Conversion of C3 to C3MA in the presence of hC3Nb1 was evaluated using cation exchange chromatography. A solution of 0.8 mg/mL of native C3 was dialyzed into 30 mM Na_2_HPO_4_ pH 8.0 and incubated with and without a 3-fold molar excess of hC3Nb1 for 10 min. After the pre-incubation, 10 mM methylamine was added, and the mixture incubated at 37 °C. At timepoints 0, 30, 60, 120, 180 min, 50 µg of reacted C3 was diluted 20-fold with equilibration buffer containing 10 mM MES pH 6.0, 50 mM NaCl buffer and loaded into a Source 15S 1 mL equilibrated in the same buffer. The applied sample was eluted with a linear gradient from 50 to 375 mM NaCl in a 20 mL gradient. The C3, C3* and C3MA species were quantified by integration and deconvolution of the elution profiles obtained in the presence and absence of hC3Nb1. For this analysis, the hC3Nb1 and C3MA complex was considered to be formed with a 1:1 stoichiometry.

### Methylamine treatment of pro-C3 and FI degradation of pro-C3 and C3ΔTE

Pro-C3 was treated with 200 mM MA as described for native C3 and applied to the 1 mL Mono S column for analysis. To analyze FI cleavage of C3b, pro-C3, pro-C3*, and C3MA, proteins were pH adjusted with HEPES pH 7.5 before being mixed with the recombinant cofactor fragment CR1 (8-10) ^52^ at a 1:400 ratio (C3b,C3MA and C3ΔTE) or 1:75 ratio (pro-C3 and pro-C3*) and with FI (Complement Technology) at a 1:100 ratio (C3b,C3MA and C3ΔTE) or 1:20 ratio (pro-C3 and pro-C3*).

### Negative stain EM of pro-C3 and C3ΔTE

Copper grids (G400-C3, Gilder) covered with amorphous carbon by evaporation on collodion were glow-discharged for 45 s at 25 mA using an easiGlow instrument (PELCO). 3 μl of pro-C3 at 1.8 μg/mL was incubated on the grid for 1 min before blotting. The grid was washed with 3 μl of 2% (w/v) uranyl formate for 15 s, followed by blotting and incubation with another 3 μL of uranyl formate for 1 min. The grid was allowed to dry before imaging in a 120 kV Tecnai G2 Spirit electron microscope. Automated data collection was done using SerialEM ^53^ at a nominal magnification of 67k and a defocus ranging from 1.5 to 2.5 μm. Using cryoSPARC ^54^, 31 exposures were used to pick 1,264 particles by picking against 130-150 Å Gaussians, which were filtered by a round of 2D classification down to 782 particles. Multiple 3D ab-initio reconstructions were made from this particle stack, one of which had 88% of the particles assigned, and thus this was used for a single round of homogenous refinement.

Grids for nsEM with C3ΔTE was prepared and imaged identically, except that the sample was added at a concentration of 1.5 μg/mL and the nominal defocus was constantly 1.4 μm. Processing was also done identically to pro-C3, and 19k particles were picked from 101 exposures. After 2D classification, 14k particles remained, 7.8k of which were assigned to the same class and subsequently refined by homogeneous refinement.

### Bio-layer interferometry with the CR3 headpiece

BLI experiments were carried out using an Octet Red96 (ForteBio, Fremont, USA) operated at 30 °C with shaking at 1000 rpm. The running buffer for all experiments was 20 mM HEPES pH 7.5, 150 mM NaCl, 5 mM MgCl_2_, 1 mM CaCl_2_ supplemented with 1 mg/mL bovine serum albumin. After dissociation, the streptavidin biosensors (Fortebio) were regenerated with 20 mM HEPES pH 7.5, 1 M NaCl, 50 mM EDTA. For loading, the streptavidin biosensors were dipped in 10 µg/mL biotinylated iC3b. A competition assay was performed to examine the binding of the CR3HP to iC3b in the presence of C3b, C3MA and iC3b. In this experiment, the association of 50 nM CR3 alone or mixed with either 10-fold molar excess of C3b, C3MA or iC3b was measured for 180 s followed by 180 s dissociation.

### Cryo-EM structure determination of C3MA

The C3MA:hC3Nb1-N108C dimer was prepared by mixing C3MA with the hC3Nb1 N108C dimer in a 1.1:0.5 molar ratio. The sample was run on a Superdex 200 increase 3.2/300 column (column) equilibrated in 20 mM HEPES pH 7.5, 150 mM NaCl at RT. Purified C3MA-hC3Nb1 dimer (3.5 μL at 0.6 mg/mL) was applied to holey carbon grids C-flat 1.2/1.3 300 mesh (Protochips) previously glow-discharged for 45 s at 15 mA using a GloQube (Quorum Technologies). The grids were vitrified on a Mark IV Vitrobot (Thermo Fisher Scientific) at 4 °C and 100% relative humidity with a blot force of 1 and blot time of 4.5 s. Data were collected on a Titan Krios G3i instrument with an X-FEG operated at 300 kV and equipped with a Gatan K3 camera and a Bioquantum energy filter (Gatan) using a slit width of 20 eV. In total, 6,093 movies were collected using aberration free image shift data collection in EPU (Thermo Fisher Scientific) as 1.5 second exposures in super-resolution mode at a physical pixel size of 0.647 Å/pixel with a total electron dose of 59.7 e^-^/Å^2^ fractionated over 56 frames.

Processing was performed in cryoSPARC v3.3.1 ^54^ as summarized in Supporting Figure 2. Patch Motion Correction and Patch CTF were performed before micrograph curation based on CFT estimations, defocus, total frame motion and ice thickness. Particles were picked with a template picker as implemented in cryoSPARC v3.3.1. Templates were generated from a 3D volume reconstructed from a blob picked particle stack curated by 2D classification and heterogeneous refinement. Template picked particles were 4x downsampled and further sorted by 2D classification and two rounds of heterogeneous refinement. Homogenous refinement of the resulting 834,749 particles indicated remaining heterogeneity, which was resolved using 3D variability analysis ^55^ to a subset containing 197,438 particles (Class 1, Supporting Figure 2D). Homogenous refinement and local refinement were done using a dilated mask encompassing the entire C3MA-nanobody dimer.

A model was built using the monomeric C3b-hC3Nb1 nanobody complex (PDB entry 6ehg) as a starting model. ChimeraX ^56^ was used for map inspection and initial fitting of models into 3D reconstructions. Programs from the PHENIX package ^57^ were used for real-space refinement, analysis and validation of cryo-EM structures. The program Coot ^58^ was used for manual rebuilding of models prior to real-space refinement in PHENIX. Model validation with MolProbity as implemented in PHENIX with relevant refinement and validation metrics is presented in Supporting Table 2. The map used for docking of ANA was obtained by blurring the 2.9 Å non-sharpened map of the C3MA-nanobody dimer with a B-factor of 133 Å^2^ in Coot.

### Cryo-EM structure determination of C3* and the C3*-hC3Nb2 complex

C3* was obtained by incubating C3 at 37 °C with 200 mM methylamine pH 8.5 for 15 minutes before moving the sample on ice and purifying it as was done for the time-course experiments. Gold grids Au-Flat 0.6/1.0 300 mesh (Protochips) were glow-discharged for 45 s at 15 mA using a GloQube instrument (Quorum Technologies). Blotting of 3.0 μL C3* sample at 0.2 mg/mL for 6 s (untilted) or 3 s (25° tilt) was done in a Mark IV Vitrobot at 4 °C and 100% relative humidity at a blot force of –10. Data collection was done in EPU 3.2 on a Titan Krios G3i electron microscope operated in nanoprobe mode at 300 kV with a defocus between 0.7 μm and 2.4 μm onto a K3 detector with a Bioquantum energy filter (20 eV slit width), a C2 aperture of 50 μM, spot size 3 and a beam diameter of 0.96 μm (untilted) or 0.92 μm (25° tilt). Images were gain-corrected and 2x binned continuously, resulting in 7,366 (untilted) and 897 (25° tilt) movies with a calibrated pixel size of 0.647 Å/px. A total dose of 59.5 e^-^/Å^2^ (untilted) or 60.3 e^-^/Å^2^ (25° tilt) was split into 53 fractions with a total exposure time of 1.4 s.

Image analysis was performed in cryoSPARC v4. The movies were patch motion-corrected and patch CTF-estimated, and the untilted micrographs were filtered based on ice-thickness and CTF-fit. Initial picks were obtained by template picking against a 30 Å low-passed reconstruction from a screening dataset, which was in turn determined by picking against 100-150 Å diameter elliptical gaussian blobs. After several rounds of 2D Classification, 1000 micrographs with 19,310 particles were used to train a ResNet8 model in Topaz ^59^, which was subsequently used to pick from all 6,689 micrographs, and particle images were extracted in 300 Å boxes and downsampled by a factor 4. The particles were filtered based on picking score, 2D Classification, Ab-Initio Reconstruction into 6 classes and several rounds of Heterogeneous Refinement. A consensus map was obtained by refinement of the cleaned stack of 92,325 particles in steps of maximum alignment resolution using Homogeneous Refinement and, subsequently, Non-Uniform Refinement with a maximum alignment resolution of 12 Å and dynamic masking at a threshold of 0.4 with 6 Å mask dilation and padding to 14 Å. Anisotropy estimation using CryoEF ^60^ was used to determine that a stage tilt of 25° was optimal for further data collection to avoid orientational bias. The consensus map low-passed to 20 Å was used to template-pick the tilted micrographs, and 204,702 particles were extracted as before. After a round of 2D Classification, Ab-Initio Reconstruction and Heterogeneous Refinement, 65,247 tilted particles were combined with the 92,325 particles used for the consensus map along with 48,324 other untilted particles and aligned against the consensus map in a Non-Uniform Refinement with scale minimization at each iteration. Over several iterations of Local Refinement with a focused map on the MG-ring with decreasing prior standard deviations and 3D Classification, the final stack of 56,532 particles was obtained. The particles were re-extracted with the same physical box size but downsampled by a factor of 3 and used for a final Local Refinement against the map focused on the MG-ring. An ab-initio reconstruction showing the TE and CUB domains was used for Homogenous and Local Refinement and a reconstruction centered on MG7 and showing C345c was used for Homogenous Refinement yielding the final consensus map, both with the same particle stack. The maps were analyzed for heterogeneity and anisotropy by Local Resolution Estimation in cryoSPARC and 3DFSC ^61^ on the web-server (https://3dfsc.salk.edu/).

The three focused maps were aligned using the “Fit in map” tool in ChimeraX ^56^. Domains from C3 (PDB entry 2a73) were placed by hand, linking residues modelled in Coot ^58^, and the model was used to combine the maps using phenix.combine_focused_maps ^57^. The model was then real-space refined against the combined map using data to 8.1 Å, and the refined model was used to generate a new combined map. The Asn939 glycosylation was added to the model from C3b (PDB entry 5fo7), and a final round of refinement was performed.

C3*-hC3Nb2 was prepared by adding hC3Nb2 to C3* purified as described above in a 1:1 ratio. Grids were prepared similarly to C3*, except that a blotting time of 7 s and blot force of -8 was used with grids for untilted collection, while an UltrAuFoil 0.6/1.0 300 mesh (Quantifoil) grid was used with the C3*-hC3Nb2 sample at 0.3 mg/mL and blotted for 11 s at blot force -8 for tilted collection at an angle of 20°. Movies were collected at defoci between 0.6 μm and 2.3 μm, the beam diameter was 1.16 μm, and the total dose was 59.3 e^-^/Å^2^ (untilted) and 59.7 e^-^/Å^2^ (20° tilt). Image analysis was also performed similarly to C3*.

Particles were picked against 130-150 Å diameter gaussian elliptical blobs, extracted in 464 px boxes and downsampled to 156 px. The 142k untilted particles were filtered by a single round of 2D Classification. 56k of these were used for Ab-Initio Reconstruction into 10 classes, which were further used for Heterogeneous Refinement. Of these, 8 classes (48k particles) showed a recognisable part of C3* and were aligned by Non-Uniform Refinement against an ab-initio volume showing the full MG-ring. Tilted particles were picked and extracted identically, and the 87k tilted particles were pooled with the 48k untilted particles and used for Ab-Initio Reconstruction and Heterogeneous Refinement into 6 classes. The 44k particles assigned to classes showing features of C3* were used for a single round of Heterogeneous Refinement using two volumes showing the MG-ring. The class reaching the highest resolution (22k particles) was refined further by Non-Uniform Refinement. Map anisotropy was assessed by Orientation Diagnostics and the median, local resolution at FSC=0.5 was found to be 5.1 Å by Local Resolution Estimation.

### C3* SAXS data collection

Static SAXS data for C3* was measured at the P12 beamline at Petra III (DESY, Hamburg, Germany). C3* was prepared and purified on-site as described above, concentrated at 4 °C in the elution buffer of 20 mM MES pH 6.0, 200 mM NaCl to 1.25 mg/mL and measured within 4 hours of the end of the incubation with methylamine. The scattering data was analysed using Primus in the ATSAS package ^62^. As the Guinier plot only increased non-linearly for the points with the two lowest **s**, this was most likely due to incomplete blocking of the direct beam and aggregation was negligible. However, estimation of the molecular weight in Primus showed a credibility interval of [195;264] kDa, suggesting some degree of dimerization as observed for native C3 and C3MA at lower concentrations of NaCl ^63^. This was also apparent from the R_max_ of 21.8 nm from the distance distribution function. To reduce the signal from putative dimers of C3*, data at **s** < 0.34 nm^-1^, corresponding to a D_max_ of 18.5 nm were omitted for further analysis. The truncated data (0.34 nm^-1^ < s < 3.0 nm^-1^) suggested a molecular weight credibility interval of [177;221] kDa using the 30 remaining points with smallest **s**, and an R_max_ of 15.1 nm was reached from the distance distribution function. A model was fitted to the truncated data using CORAL ^62^ by fixing the C3* model from cryo-EM and allowing placement of MG3, ANA and high-mannose glycosylations on Ans85 and Asn939 ^64^. The C1 atom of the Asn-linked NAG residues were restrained to be 6 Å from the Cα atom of the connected asparagine. The ANA - (Nt-α) link which was restrained to 15 Å between Ala747 and Glu753. Using CRYSOL, structures of native C3 (PDB: 2a73), C3MA (monomer from our cryo-EM structure), C3* cryo-EM model missing MG3 and ANA, C3* cryo-EM model with MG3 and ANA placed as in native C3, and the rigid-body fitted model were evaluated against the either the full range of data (**s** < 3.0 nm^-1^) or the truncated data (0.34 nm^-1^ < **s** < 3.0 nm^-1^). Model distance distribution functions were obtained by calculating the theoretical scattering curve of atomic models using CRYSOL as above and then calculating the distance distribution function using GNOM.

### Structural analysis and comparisons

All figures were prepared with PyMOL or ChimeraX. To obtain an improved model of native bovine C3, a starting model was obtained by combination of the deposited PDB entry 2B39 ^15^ and the Alphafold2 model of bovine C3. The model was rebuilt in Coot and refined and validated in PHENIX.

## Acknowledgements and funding

We are grateful for the assistance with data collection given by the staff at the Aarhus University EM facility, EMBION and we thank Jesper L Karlsen for advice on EM data processing. Søren Vrønning Hoffmann and Nykola Jones from the ASTRID CD spectroscopy beamline assisted during data collection and analysis. We would like to thank Clement Blanchet for assistance during SAXS data collection at the EMBL P12 beamline at the PETRA III storage ring (DESY, Hamburg, Germany). This work was supported by Lundbeck Foundation BRAINSTRUC center (R155-2015-2666 to G.R.A and B.B.K), the Novo Nordisk Foundation (NNF18OC0052105, and NNF20OC0065238 to G.R.A, NNF18OC0032724 BIO-MS to J.J.E., and NNF18OC0033926 to B.B.K), and the Danish council for independent research (10.46540/2032-00111B to J.J.E).

## Author contributions

**Trine AF Gadeberg, Martin H Jørgensen, Heidi G Olesen, Josefine Lorentzen, Sean DL Harwood, Ana V Almeida**: Experimental work, data analysis, manuscript preparation. **Marlene U Fruergaard, Rasmus K Jensen, Philipp Kanis, Henrik Pedersen, Emil Tranchant, Ida Thøgersen, Steen V Pedersen**: Experimental work and data analysis. **Birthe B Kragelund, Joseph A Lyons, Jan J Enghild**: Data analysis. **Gregers R Andersen**: Experimental work, data analysis, manuscript preparation, supervision, project management.

## Data availability

The MS data files used for intact mass determination of C3a and identifying the trypsin-generated disulfide-bonded C3a tetrapeptide have been deposited to the ProteomeXchange Consortium via the PRIDE ^65^ partner repository with the dataset identifier PXD041974. Coordinates and maps for the C3MA are available as PDB entry 8oq3 and EMDB entry EMD-17103. The three partial maps for C3* are available at EMDB as entries EMD-17326, EMD-17325 and EMD-17328 along with the combined map as entry EMD-17327. The updated structure and original diffraction amplitudes for bovine C3 is available as PDB entry 8cem. The CD data is available in supporting table 7. The SAXS data is available as SASBDB entry SASDTU8.

