## Supporting information for "Cryo-EM analysis of complement C3 reveals a reversible major opening of the macroglobulin ring"

### Supporting information figure legends

**Supporting Figure 1. Sample preparation for C3MA structure determination. A**) Crystal packing dimer of the C3b-hC3Nb1 complex in PDB entry 6ehg. **B**) Top view of the nanobody dimer from panel A with the two Asn108 residues. **C**) Non-reducing SDS-PAGE analysis of the hC3Nb1 N108C mutant before and after dimerization with DTNB. The nanobody dimer migrates at 28 kDa as compared to the monomer migrating at 12 kDa. **D**) SEC purification of the 2:2 C3MA:hC3Nb1 complex for cryo-EM structure determination. The light shaded volume in fraction 7 was used for the preparation of grids. The peak at 1.25 ml contains excess C3MA. **E**) Blot used for N-terminal sequencing with two identical lanes of sample from the dark shaded fraction 9 in panel D. The band at the top starts at serine 672 (prepro numbering) confirming that the α-chain in C3MA is intact. As expected, the lower band starts at the N-terminal serine 23 in the C3MA β-chain. The weak band below the 20 kDa marker was not sequenced but is likely to correspond to the reduced monomeric hC3Nb1 N108C.

**Supporting Figure 2. Cryo-EM structure analysis of the C3MA-hC3Nb1 dimer. A**) Single particle analysis pipeline outlining the processing and refinement performed to obtain the model. **B**) Example of -1.9 µm defocus micrograph post patch motion-correction. **C**) Representative 2D class averages of C3MA from cryoSPARC. **D**) Four clusters identified by 3D variability analysis. The cluster chosen for downstream refinement is indicated. **E**) Map from homogenous refinement with C1 symmetry. **F**) Map from local refinement with C1 symmetry**.**

**Supporting Figure 3. Aspects of the C3MA structure and comparison of C3b with C3MA**. **A**) The 3D reconstruction of the C3MA-hC3Nb1 dimer allows an unambiguous tracing of the Nt-α' region in C3MA. **B**) The 3D reconstruction confirms the formation of the disulfide bridge between the two hC3Nb1 nanobodies at Cys108. The map is contoured at 6σ in panels A-B. **C**) The superposition of the C3MA and C3b (PDB entry 6ehg) structures on their MG1-8 domains illustrates their close resemblance. The structures superimpose with a root mean square deviation of 0.66 Å over 897 $C_\alpha$ atoms. **D)** Magnified view comparing the MG-ring conformation in C3MA (colored by domain) and C3b (grey). **E**) Bio-layer interferometry confirms the functional similarity of C3b and C3MA. Unlike fluid phase iC3b, neither C3b nor C3MA are able to compete with sensor-bound iC3b for binding of fluid phase complement receptor 3 (CR3). A 10-fold molar excess of iC3b, C3MA and C3b was used relative to the CR3 headpiece in the fluid phase. The BLI experiment was conducted in two biological replicates.

**Supporting Figure 4. MS analysis of C3a generated from native C3 and C3MA. A**) Time course of trypsin digestion comparing the proteolytic susceptibility of C3 and C3MA, the latter more readily digested in accordance with the ANA domain being dynamically attached to the rest of the molecule. The bands between 150-200 kDa contain residual non-reduced C3 and C3b. **B-C**) MS analysis of intact C3a$_T$ from C3 and C3MA. The total ion chromatogram, average MS1 spectrum of all peptides, and deconvoluted masses from the average MS1 spectrum are shown. Masses matching C3a-like peptides are annotated A, B, C, D, and E, with 0-3 methionine oxidations. The sequences of these peptides are given in Table S3. **D-E**) Disulfide investigation in C3a$_T$ from C3 or C3MA. The peptides from low-pH (pH 6) trypsin digestion of C3a$_T$ were separated by RP-HPLC, and the fraction containing "peak 1" was determined by Edman sequencing to contain the disulfide-covering peptides. LC-MS/MS analysis of this fraction from each C3 preparation produced an MS2 spectrum for a tetrapeptide consistent with the known disulfide pattern of C3a. These MS2 fragments were insufficient for de novo disulfide mapping of C3a but allowed only for disulfide scrambling between adjacent cysteines. Furthermore, the high degree of similarity between the two MS2 spectra indicates that the disulfide pattern is identical in each sample. **F)** C3MA is formed from C3* in the absence of MA at pH 6 during long-term incubation on ice (top) and further stimulated by short-term incubation at 37° C (bottom). This argues against disulfide shuffling being essential for the reaction. The experiments presented in this figure were conducted one time.

**Supporting Figure 5. Details of SRCD analysis of multiple functional states of C3. A**) Time course conversion of C3 to C3MA in the presence of 10 mM methylamine measured by Synchrotron Radiation Circular Dichroism (spectra as black dots) and secondary structure deconvolution by DicroWeb (red line) showing a practically constant secondary structure content throughout the reaction. **B)** The reaction progression was monitored in parallel by ion exchange chromatography. **C)** C3, C3* and C3MA distribution throughout the reaction obtained from deconvolution of the chromatograms obtained as illustrated in panel B. **D)** Secondary structure content for each time point generated with the Dichroweb analysis tool. The 10 best results for the secondary structure prediction were averaged and 95% confidence intervals were obtained. The measurements presented in the figure were conducted one time for each time point.

**Supporting Figure 6. Details of cryo-EM analysis of C3*. A)** Elution traces from a 1 ml Mono-S column of C3 incubated with 200 mM methylamine pH 8.5 at 37 °C for the designated time periods. Distinct peaks are seen for C3, C3* and C3MA. **B)** A representative micrograph at -1.5 µm defocus after patch motion-correction obtained from a Mono-S C3* peak sample under the conditions in panel A but after 15 minutes of incubation. **C)** Workflow showing the steps of C3* cryo-EM SPA with annotation of the kept particles at each step. The final, reported resolutions are the reported global resolutions from the final refinement, except for the consensus map which was estimated using the median local resolution at FSC = 0.5. **D)**

Selection of 2D classes obtained from the untilted data. For one class, the area where the MG3 domain is clearly missing is designated by a red arrow.

**Supporting Figure 7. SAXS analysis of C3*. A)** Raw scattering curve for C3*. **B)** Distance distribution function calculated from the full range data. **C)** Theoretical distance distribution functions for native C3 (PDB: 2a73), the C3MA cryo-EM model, and the C3* cryo-EM model missing MG3 and ANA. Until $r > 11$ nm, the distance distribution calculated from C3* fits the experimental P(r) much better than the distance distributions calculated from C3 and C3MA models. At longer distances, the experimental curve is likely to be dominated by the presence of dimers. **D)** Fits of theoretical scattering curves of native C3, C3MA and the C3* cryo-EM model shows the most significant differences for $0.5\ nm^{-1} < \mathbf{s} < 1.8\ nm^{-1}$, in which range the C3* cryo-EM model fits the experimental data better than any of the other two models. **E)** Model from CORAL obtained by rigid-body fitting MG3, ANA and the two glycans onto the fixed C3* cryo-EM model. **F)** Quality of fits against either the full data ($0 < \mathbf{s} < 3\ nm^{-1}$) or data truncated to $0.34\ nm^{-1} < \mathbf{s} < 3\ nm^{-1}$ of either native C3, C3MA, the C3* cryo-EM model, the C3* cryo-EM model with MG3 and ANA placed as in native C3, and the fitted model by placing MG3, ANA and the Asn939 and Asn85 glycans freely on the fixed C3* cryo-EM model.

**Supporting Figure 8. Negative stain EM confirms that pro-C3 adopts a C3-like structure. A)** A representative micrograph and selected 2D classes (left) along with the structure of native C3 (pdb entry 6ru5) docked in the 3D reconstruction obtained by negative stain EM (right). B) Illustration of why the three-helix bundle conformation is likely to occur in pro-C3 (model to the left) whereas in the experimental structure of C3, ANA is in the four-helix bundle configuration (right, pdb entry 6ru5).

**Supporting Figure 9. The MG-ring of C3ΔTE adopts a C3MA like structure. A)** SEC elution profile of C3ΔTE showing a monodisperse peak. **B)** Degradation of C3b and C3ΔTE with FI in the presence of CR1 as cofactor. Notice that the C3ΔTE $\alpha$-chain is degraded over time indicating a C3b like conformation of the MG-ring with the CUB domain juxtaposed to the MG2 domain. The experiment was repeated twice. **C)** Representative negative stain electron micrograph of C3ΔTE. **D)** Selected negative stain 2D classes. **E)** Negative stain 3D reconstruction with C3b model (PDB: 5fo7) with the TE domain removed and the CUB domain rotated to fit best into the envelope. The envelope covers the CUB domain poorly, suggesting that the domain can assume multiple orientations in relation to the MG-ring.

**Supporting Figure 10. The hC3Nb2 nanobody inhibits C3MA formation. A-C)** Time-series of C3 conversion to C3MA at 37 °C in a buffer with 10 mM methylamine pH 8.0. Examples of mass photometry profiles obtained at time-points 0 h (panel A), 3.5 h (panel B) and 8 h (panel C) by adding a two-fold molar excess of EWEµH and measuring the molecular weight distribution in the sample for 1 minute. The 15 kDa hC3Nb2 forms complexes with C3, C3* and C3MA (all 185 kDa), whereas the 50 kDa EWEµH can only form a complex with C3MA. Differences in measured, absolute masses at different time-points are due to microscope contrast drift as all measurements use the same, combined mass calibration, which fits best centrally in the time-series. **D-F)** The same reaction was followed in parallel by cation exchange chromatography (in the absence of EWEµH) for the same time-points. The excess hC3Nb2 fraction is constant during the reaction while the C3 is converted to C3MA with C3* as an intermediate. The elution profile (black dots) was fitted (red line) with a sum of Gaussian curves corresponding to each species: hC3Nb2 nanobody, C3/C3* and C3MA (black lines). Two biological replicates were conducted with mass-photometry and with ion exchange chromatography in parallel done for one of the replicates.

**Supporting Figure 11. Cryo-EM analysis of the C3*-hC3Nb2 complex. A)** Workflow representing the steps of C3*-Nb2 cryo-EM SPA with the number of kept particles at each step indicated. The final, reported resolution is the median, local resolution at FSC = 0.5. **B)** Selection of 2D classes obtained from the untilted data. For all averages showing the MG-ring, a full ring is observed. In particular, 2D class projection is now visible for the MG3 domain (blue arrow). **C)** Global FSC curves for the final map showing the C3*-hC3Nb2 MG-ring determined after mask correction in cryoSPARC, either within a tight mask, a loose mask or unmasked. The histogram of resolutions with directional FSC = 0.143 as determined using 3DFSC with the tight mask is superimposed. **D)** 3D reconstruction of the MG-ring in C3*-hC3Nb2, now showing a complete MG-ring with extra density from hC3Nb2. **E)** Close-up of the region corresponding to MG3, MG4 and hC3Nb2 seen from the side compared to panel D. The density for the MG3 domain appears noisier than the density for the hC3Nb2 nanobody, suggesting that the MG3 domain position may vary slightly even when bound by hC3Nb2.

**Supporting Figure 12. The hC3Nb1 nanobody enhances the C3MA formation. A)** Conversion of C3 to C3MA at 37 °C in a buffer with 10 mM methylamine pH 8.0 with and without hC3Nb1 followed by cation exchange chromatography profiles obtained at time-points 0 h (top), 1 h (middle) and 3 h (bottom) in the presence of a three-fold molar excess of hC3Nb1. Notice that hC3Nb1 does not associate with C3 and C3* in the cation exchange analysis. **B)** Quantitation of the three species C3, C3* and C3MA obtained by integration and deconvolution of the combined peak containing both C3 and C3*. Two technical replications were performed.

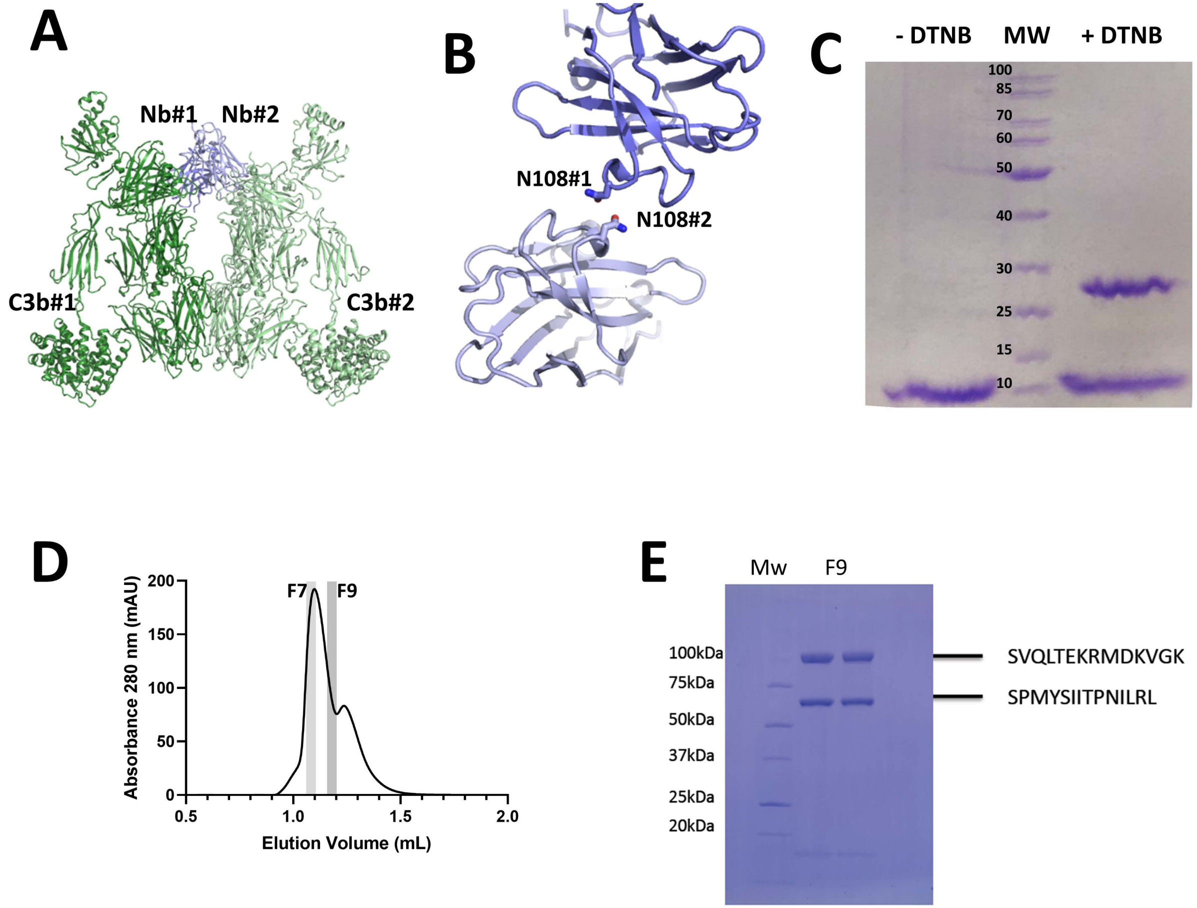
A
Nb#1
Nb#2
C3b#1
C3b#2
B
N108#1
N108#2
C
- DTNB
MW
+ DTNB
100
85
70
60
50
40
30
25
15
10
D
F7
F9
Absorbance 280 nm (mAU)
200
150
100
50
0
0.5
1.0
1.5
2.0
Elution Volume (mL)
E
Mw
F9
100kDa
75kDa
50kDa
37kDa
25kDa
20kDa
SVQLTEKRMDKVGK
SPMYSIITPNILRL

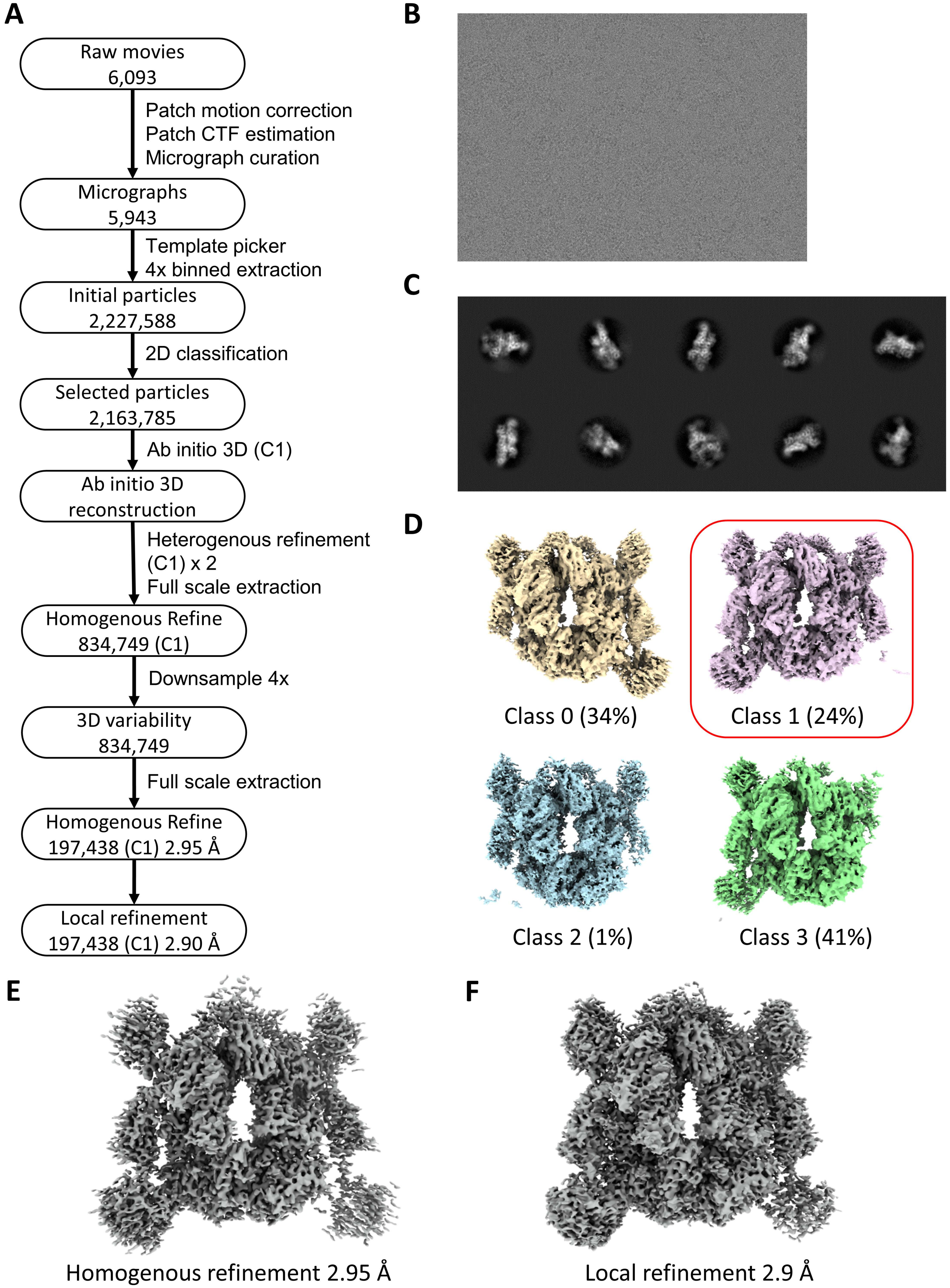

A
Raw movies
6,093
Patch motion correction
Patch CTF estimation
Micrograph curation
Micrographs
5,943
Template picker
4x binned extraction
Initial particles
2,227,588
2D classification
Selected particles
2,163,785
Ab initio 3D (C1)
Ab initio 3D
reconstruction
Heterogenous refinement
(C1) x 2
Full scale extraction
Homogenous Refine
834,749 (C1)
Downsample 4x
3D variability
834,749
Full scale extraction
Homogenous Refine
197,438 (C1) 2.95 Å
Local refinement
197,438 (C1) 2.90 Å
B
C
D
Class 0 (34%)
Class 1 (24%)
Class 2 (1%)
Class 3 (41%)
E
Homogenous refinement 2.95 Å
F
Local refinement 2.9 Å

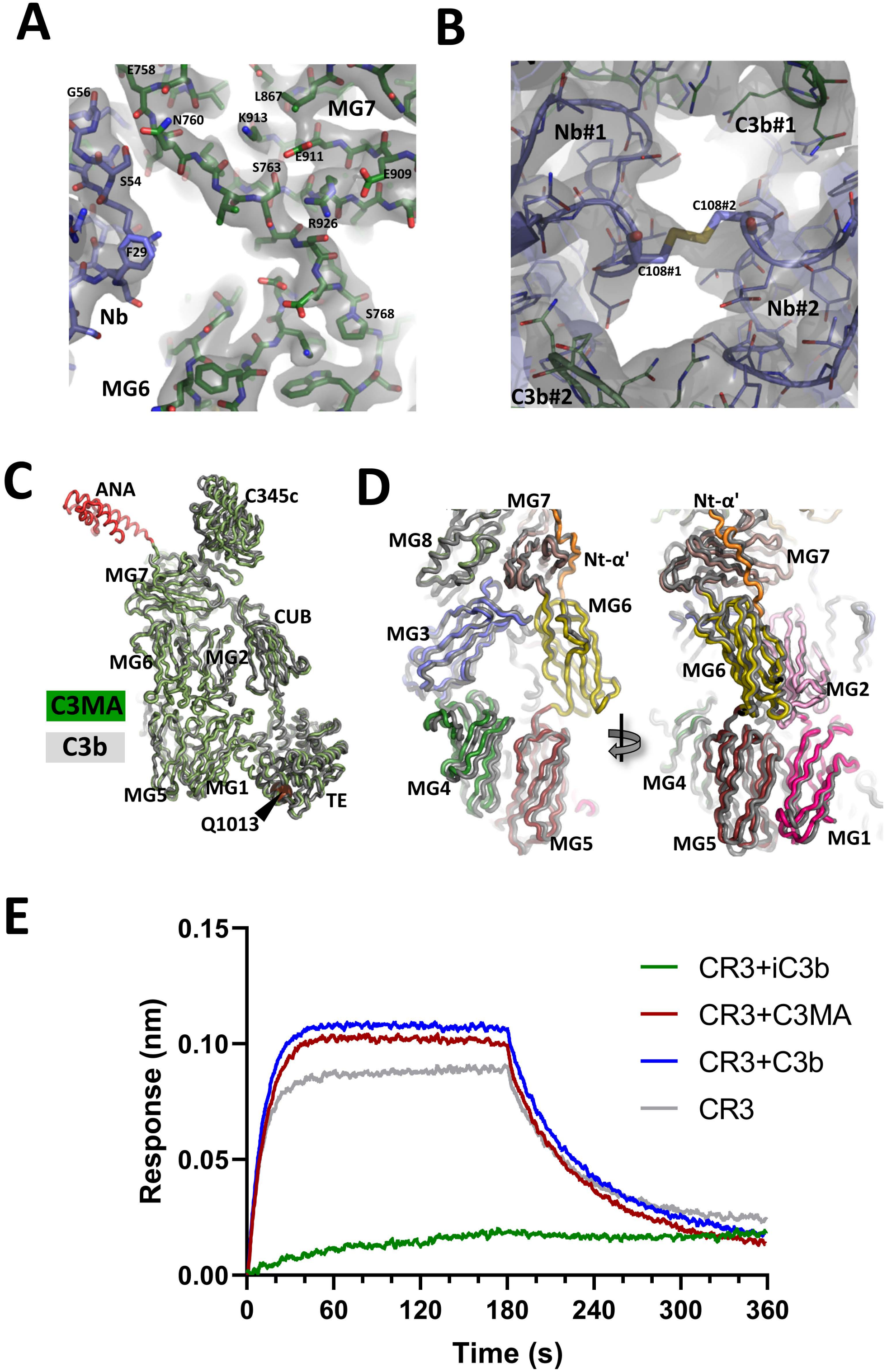

A
E758
G56
L867
MG7
N760
K913
E911
S763
E909
S54
R926
F29
Nb
S768
MG6
B
Nb#1
C3b#1
C108#2
C108#1
Nb#2
C3b#2
C
ANA
C345c
MG7
CUB
MG6
MG2
C3MA
C3b
MG5
MG1
TE
Q1013
D
MG7
MG8
Nt-α'
MG6
MG3
MG4
MG5
Nt-α'
MG7
MG6
MG2
MG4
MG5
MG1
E
CR3+iC3b
CR3+C3MA
CR3+C3b
CR3
Response (nm)
Time (s)
0.15
0.10
0.05
0.00
0
60
120
180
240
300
360

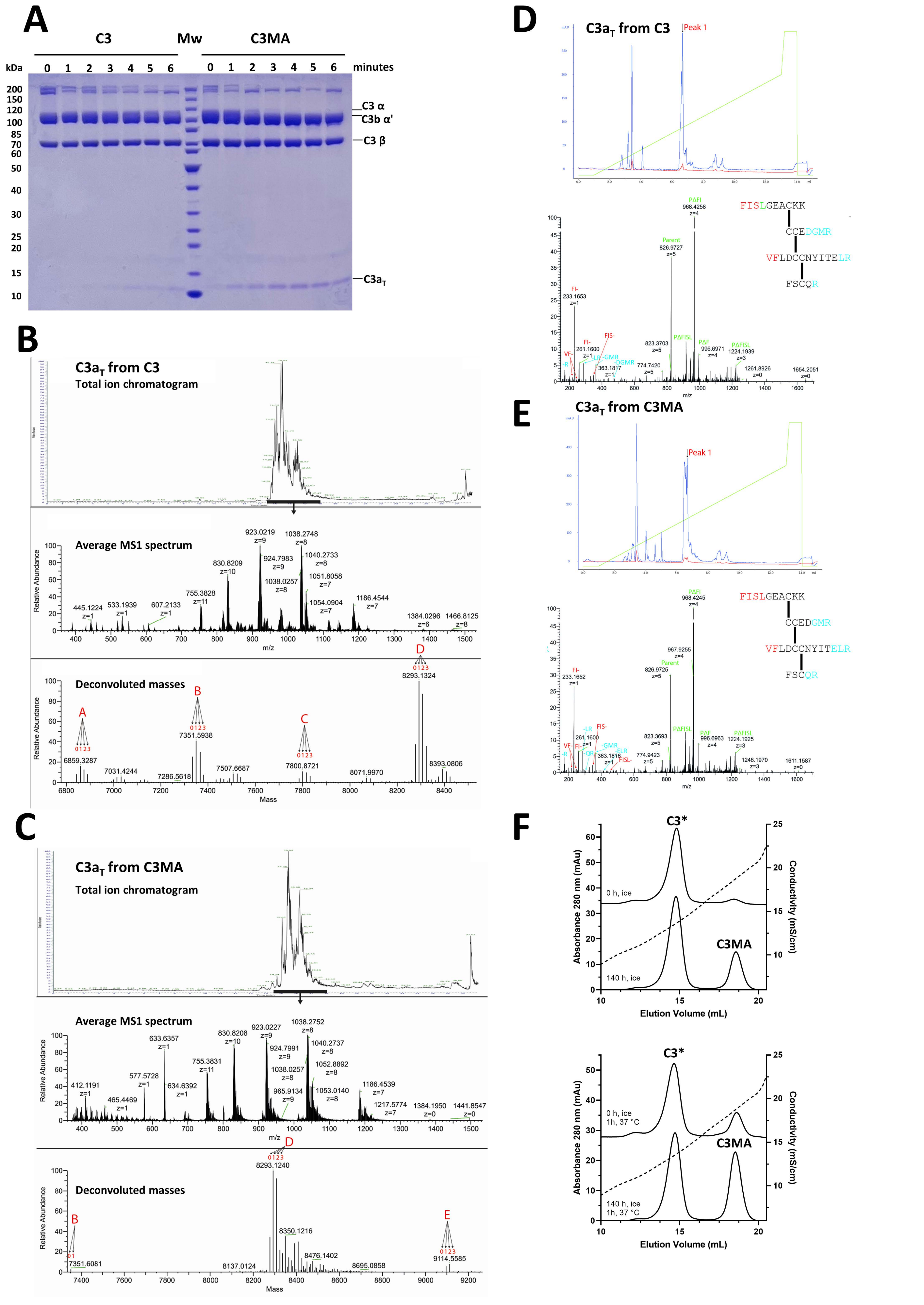

A
C3
Mw
C3MA
kDa
0 1 2 3 4 5 6
0 1 2 3 4 5 6
minutes
200
150
120
100
85
70
60
50
40
30
25
20
15
10
C3 α
C3b α'
C3 β
C3a$_T$
B
C3a$_T$ from C3
Total ion chromatogram
Average MS1 spectrum
Relative Abundance
m/z
Deconvoluted masses
Mass
6859.3287
7351.5938
7800.8721
8293.1324
8393.0806
C
C3a$_T$ from C3MA
Total ion chromatogram
Average MS1 spectrum
Deconvoluted masses
8293.1240
9114.5585
7351.6081
D
C3a$_T$ from C3
Peak 1
FISLGEACKK
CCEDGMR
VFLDCCNYITELR
FSCQR
Parent
826.9727
PΔFI
968.4258
E
C3a$_T$ from C3MA
Peak 1
Parent
826.9725
PΔFI
968.4245
F
C3*
C3MA
0 h, ice
140 h, ice
1h, 37 °C
Absorbance 280 nm (mAu)
Conductivity (mS/cm)
Elution Volume (mL)

A

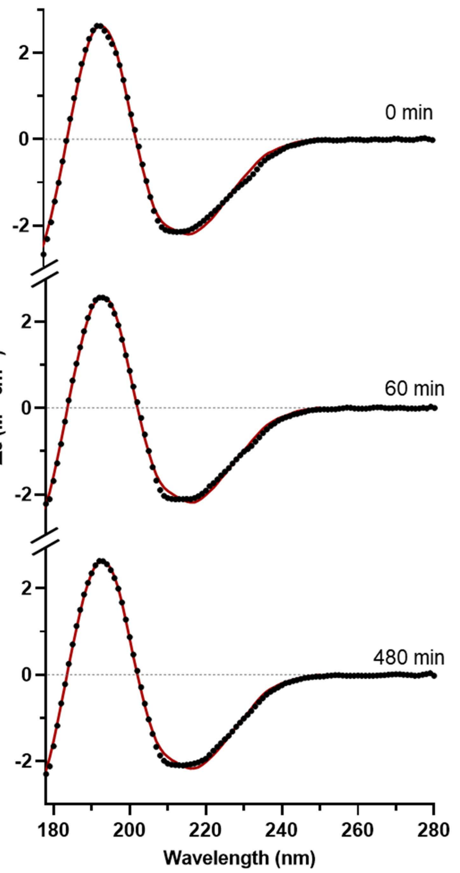


B

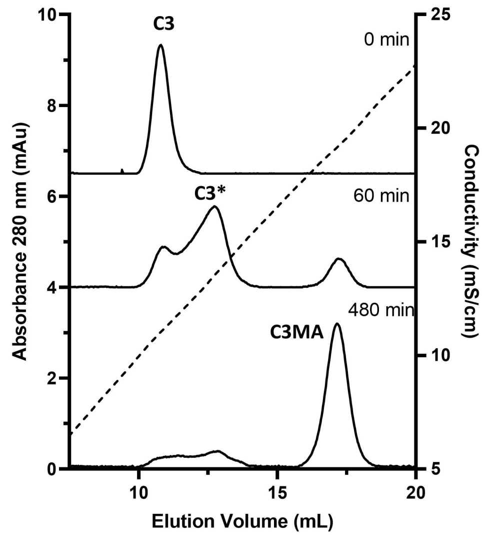


C

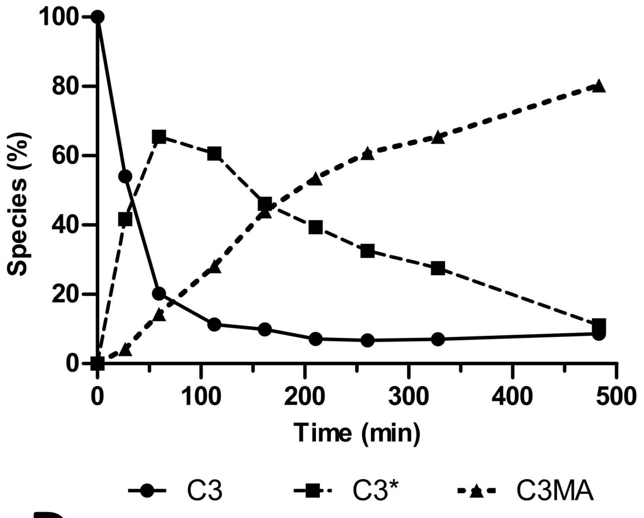


D

| | Secondary Structure Content (%) | | | |
|---|---|---|---|---|
| Time (min) | α-Helix | β-Sheet | Turns | Unordered |
| 0 | 18.9 ± 1.0 | 37.3 ± 2.1 | 10.2 ± 1.4 | 34.9 ± 2.1 |
| 25 | 17.7 ± 1.1 | 36.1 ± 2.5 | 10.6 ± 1.4 | 35.1 ± 2.2 |
| 60 | 18.2 ± 1.5 | 36.9 ± 2.1 | 10.2 ± 0.8 | 34.8 ± 2.7 |
| 110 | 18.1 ± 1.2 | 36.2 ± 3.6 | 10.2 ± 1.1 | 34.4 ± 3.1 |
| 160 | 19.1 ± 0.8 | 35.7 ± 2.5 | 11.1 ± 1.1 | 35.7 ± 2.4 |
| 210 | 17.5 ± 1.0 | 35.0 ± 2.1 | 10.7 ± 1.1 | 36.4 ± 2.0 |
| 260 | 18.9 ± 1.0 | 36.7 ± 3.0 | 11.6 ± 0.9 | 33.8 ± 3.0 |
| 330 | 18.6 ± 1.2 | 35.3 ± 3.0 | 11.3 ± 0.9 | 34.9 ± 3.0 |
| 480 | 18.0 ± 1.1 | 34.03 ± 2.5 | 11.4 ± 1.1 | 35.9 ± 2.6 |

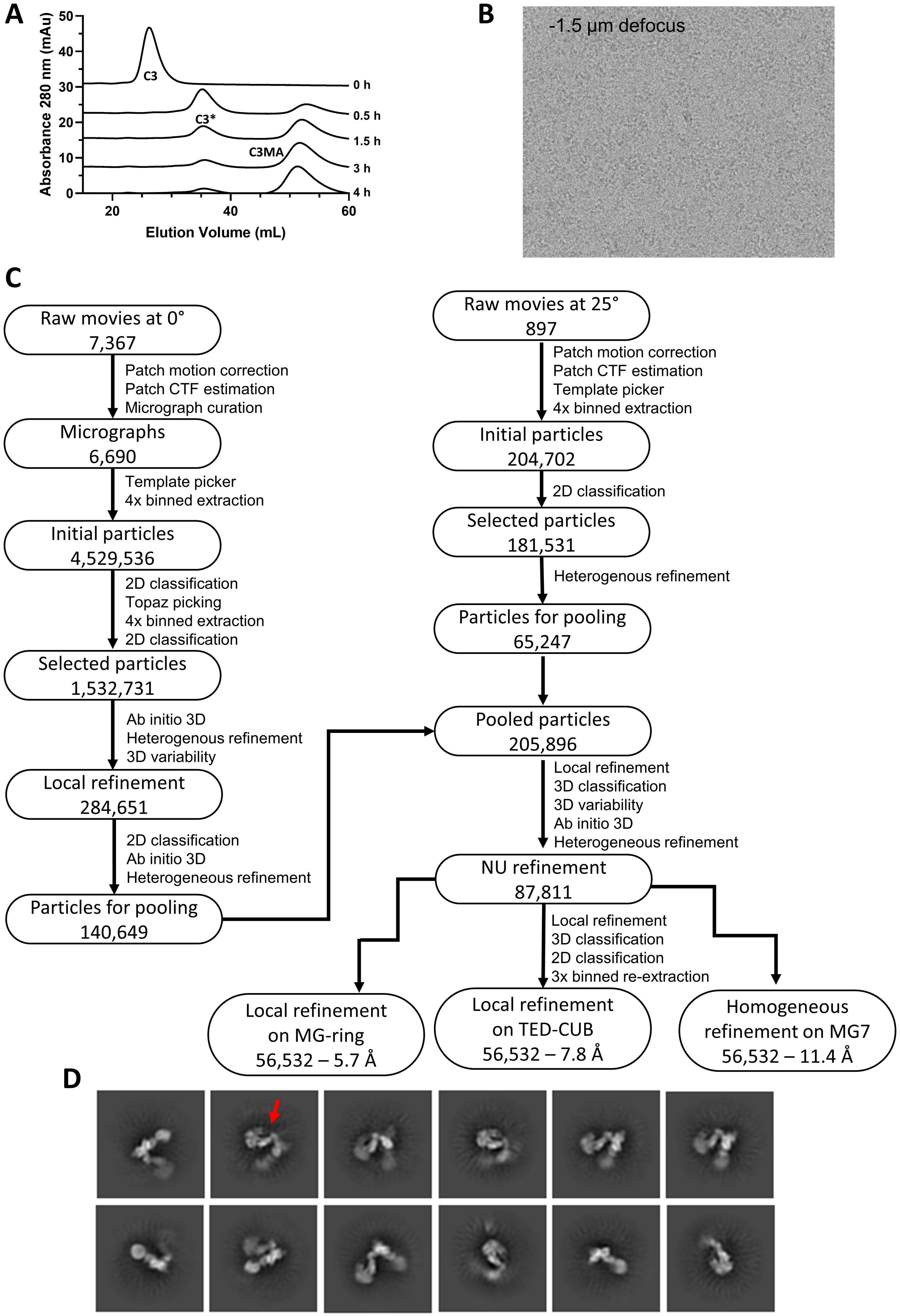
A
Absorbance 280 nm (mAu)
Elution Volume (mL)
C3
C3*
C3MA
0 h
0.5 h
1.5 h
3 h
4 h
B
-1.5 µm defocus
C
Raw movies at 0° 7,367
Patch motion correction
Patch CTF estimation
Micrograph curation
Micrographs 6,690
Template picker
4x binned extraction
Initial particles 4,529,536
2D classification
Topaz picking
4x binned extraction
2D classification
Selected particles 1,532,731
Ab initio 3D
Heterogenous refinement
3D variability
Local refinement 284,651
2D classification
Ab initio 3D
Heterogeneous refinement
Particles for pooling 140,649
Raw movies at 25° 897
Patch motion correction
Patch CTF estimation
Template picker
4x binned extraction
Initial particles 204,702
2D classification
Selected particles 181,531
Heterogenous refinement
Particles for pooling 65,247
Pooled particles 205,896
Local refinement
3D classification
3D variability
Ab initio 3D
Heterogeneous refinement
NU refinement 87,811
Local refinement
3D classification
2D classification
3x binned re-extraction
Local refinement on MG-ring 56,532 – 5.7 Å
Local refinement on TED-CUB 56,532 – 7.8 Å
Homogeneous refinement on MG7 56,532 – 11.4 Å
D

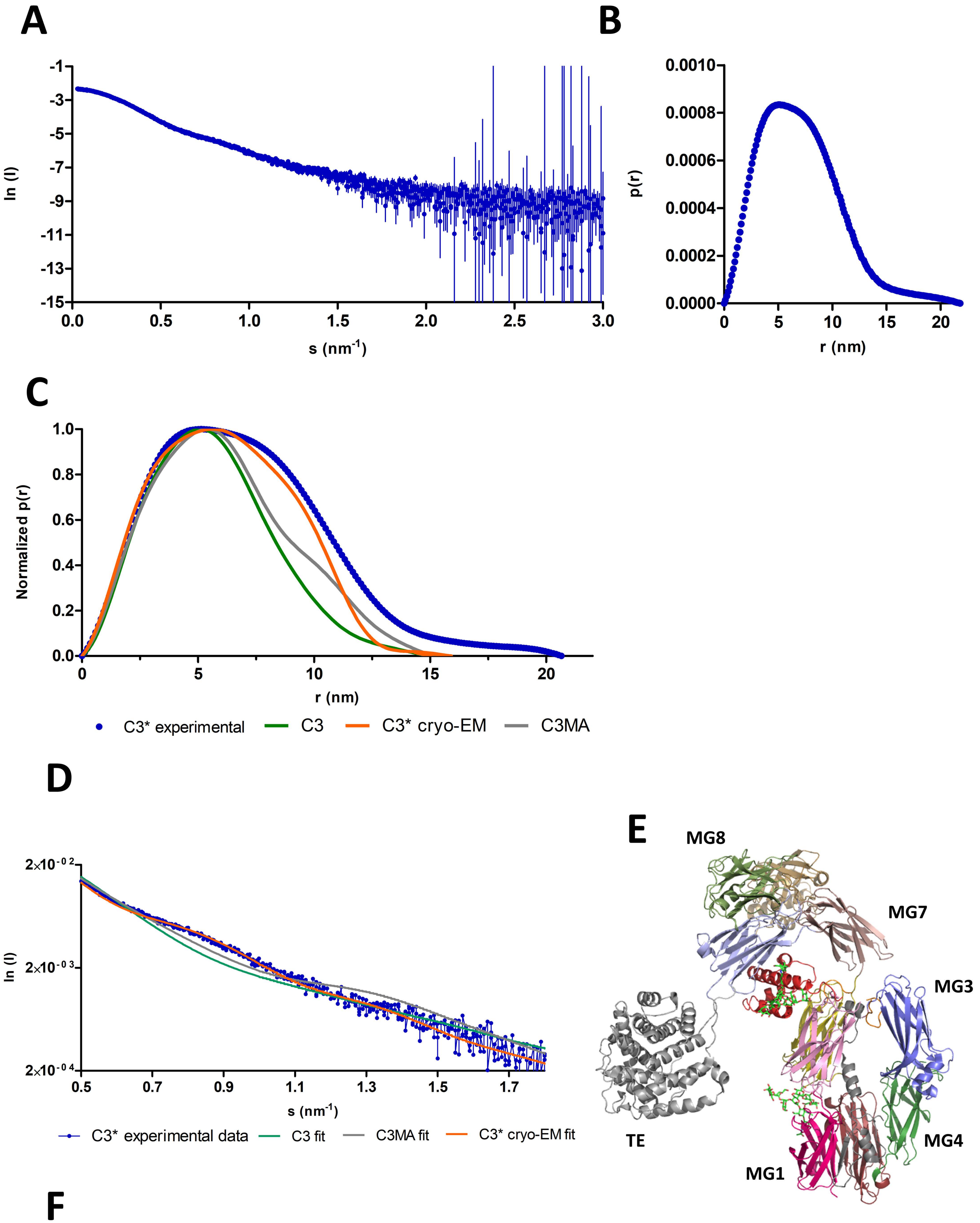


**F**

| C3* SAXS data, $\chi^2$ | C3 | C3MA | C3* cryo-EM model | C3* cryo-EM model including MG3+ANA | Rigid-body-fitted C3* with glycosylations |
|---|---|---|---|---|---|
| Full range data | 23.4 | 8.6 | 2.2 | 6.1 | 4.8 |
| Truncated data | 11.4 | 5.5 | 1.9 | 7.2 | 1.4 |

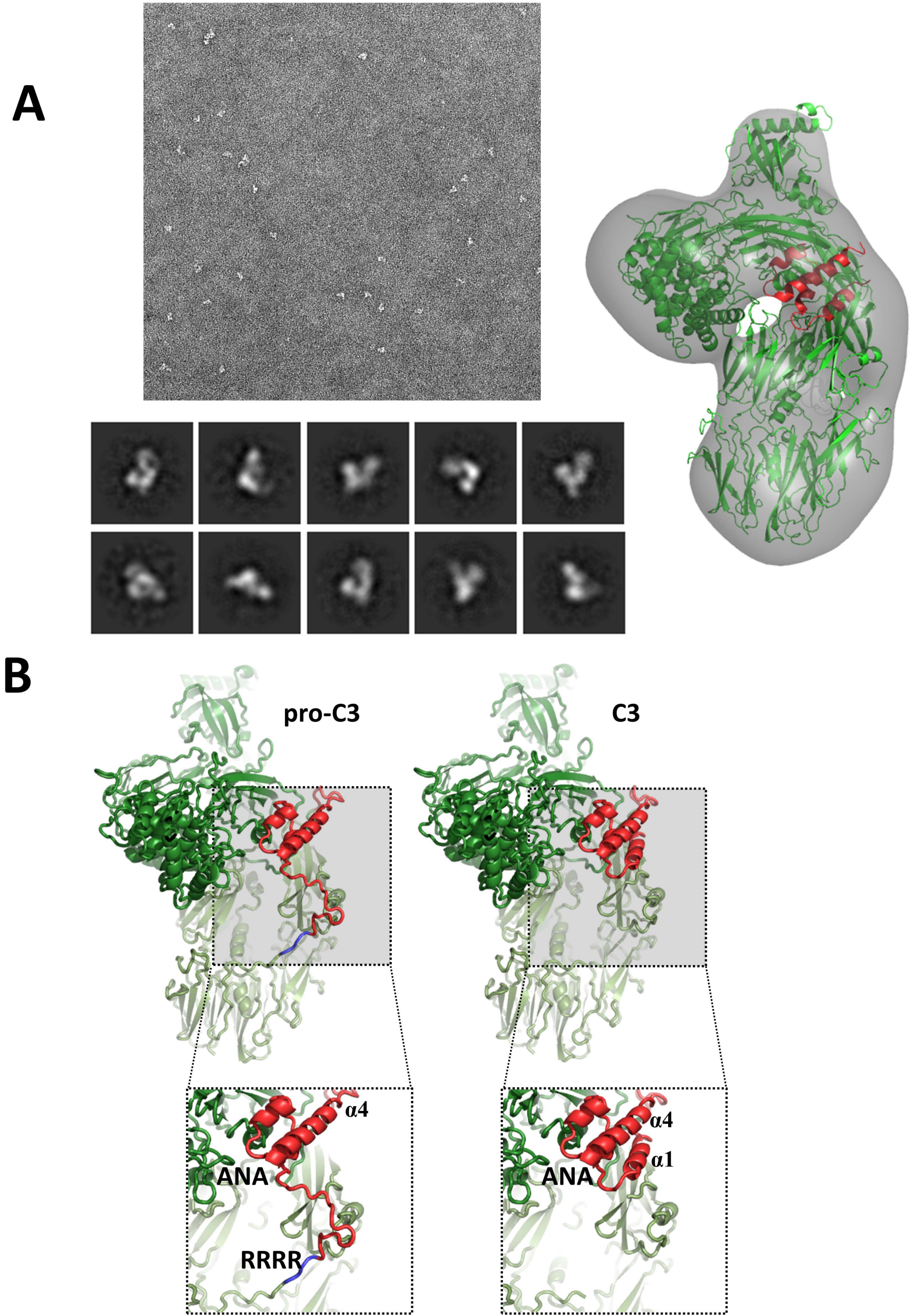
A
B
pro-C3
C3
α4
ANA
RRRR
α4
α1
ANA

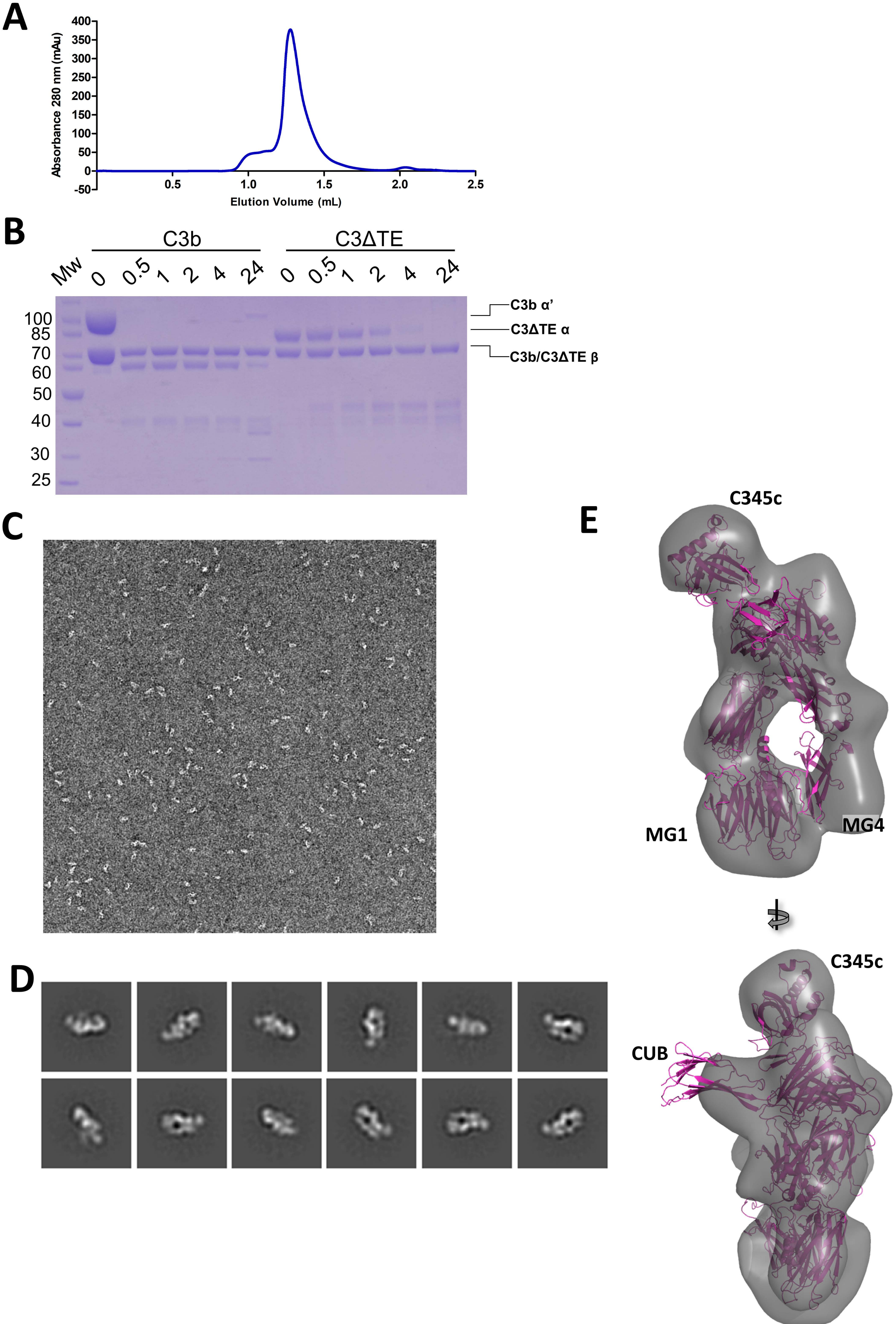
A
Absorbance 280 nm (mAu)
400
350
300
250
200
150
100
50
0
-50
0.5
1.0
1.5
2.0
2.5
Elution Volume (mL)
B
C3b
C3ΔTE
Mw
0
0.5
1
2
4
24
0
0.5
1
2
4
24
100
85
70
60
50
40
30
25
C3b α'
C3ΔTE α
C3b/C3ΔTE β
C
D
E
C345c
MG1
MG4
C345c
CUB

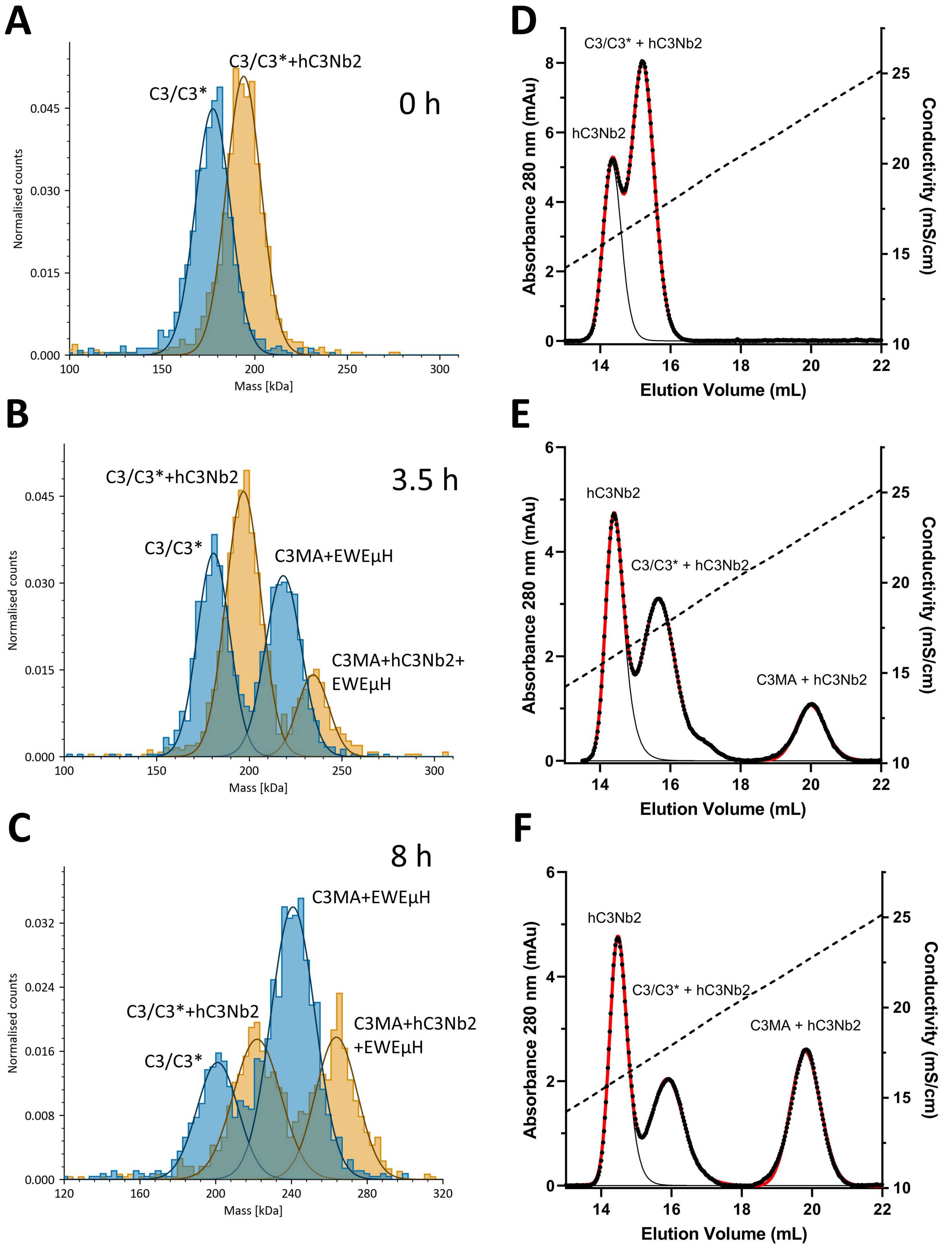

A
C3/C3*+hC3Nb2
C3/C3*
0 h
Normalised counts
Mass [kDa]
B
C3/C3*+hC3Nb2
3.5 h
C3/C3*
C3MA+EWEμH
C3MA+hC3Nb2+
EWEμH
C
8 h
C3MA+EWEμH
C3/C3*+hC3Nb2
C3MA+hC3Nb2
+EWEμH
C3/C3*
D
C3/C3* + hC3Nb2
hC3Nb2
Absorbance 280 nm (mAu)
Conductivity (mS/cm)
Elution Volume (mL)
E
hC3Nb2
C3/C3* + hC3Nb2
C3MA + hC3Nb2
F
hC3Nb2
C3/C3* + hC3Nb2
C3MA + hC3Nb2

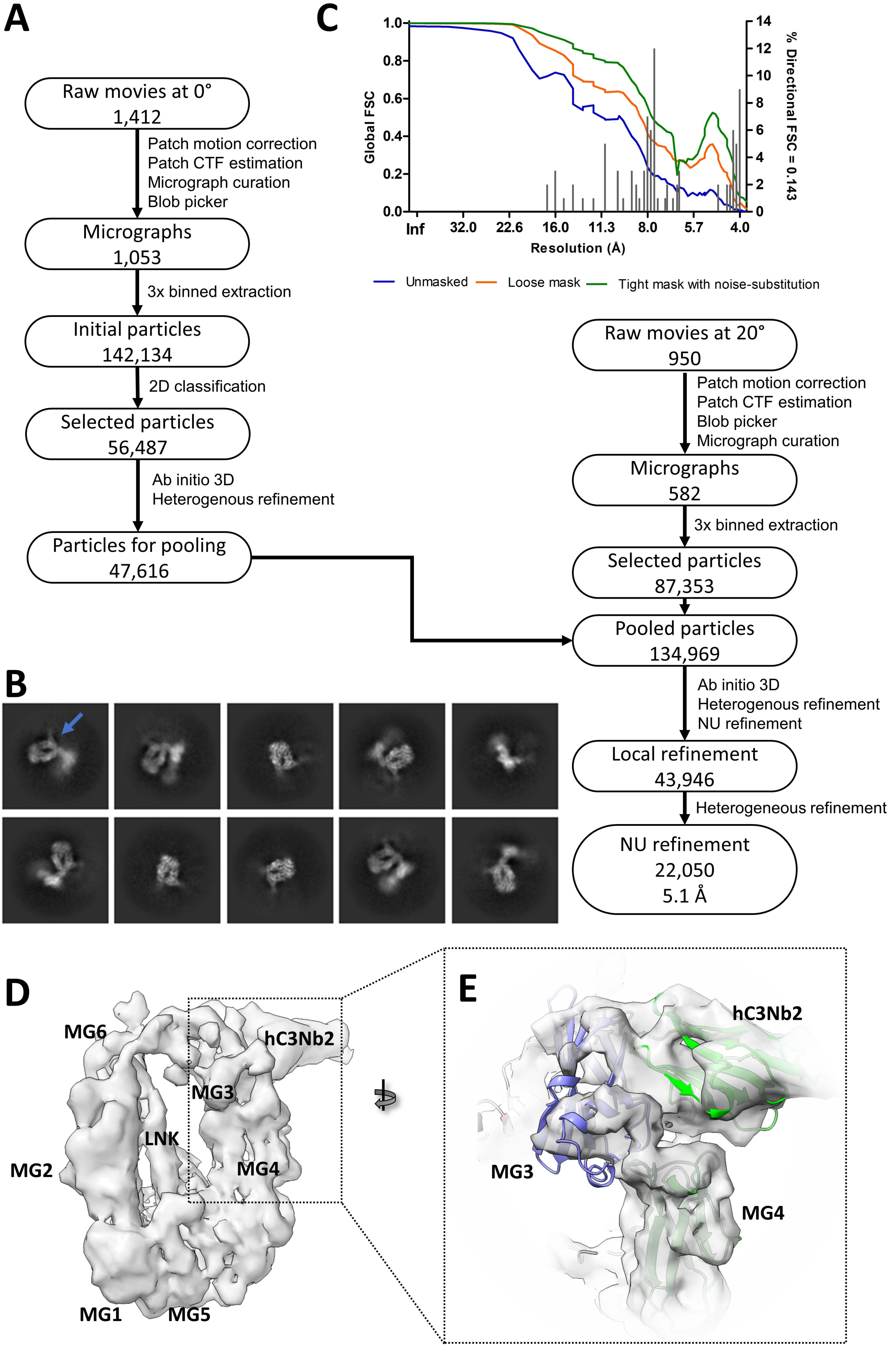
A
Raw movies at 0°
1,412
Patch motion correction
Patch CTF estimation
Micrograph curation
Blob picker
Micrographs
1,053
3x binned extraction
Initial particles
142,134
2D classification
Selected particles
56,487
Ab initio 3D
Heterogenous refinement
Particles for pooling
47,616
C
Global FSC
% Directional FSC = 0.143
Resolution (Å)
Inf
32.0
22.6
16.0
11.3
8.0
5.7
4.0
Unmasked
Loose mask
Tight mask with noise-substitution
Raw movies at 20°
950
Patch motion correction
Patch CTF estimation
Blob picker
Micrograph curation
Micrographs
582
3x binned extraction
Selected particles
87,353
Pooled particles
134,969
Ab initio 3D
Heterogenous refinement
NU refinement
Local refinement
43,946
Heterogeneous refinement
NU refinement
22,050
5.1 Å
B
D
MG6
hC3Nb2
MG3
LNK
MG2
MG4
MG1
MG5
E
hC3Nb2
MG3
MG4

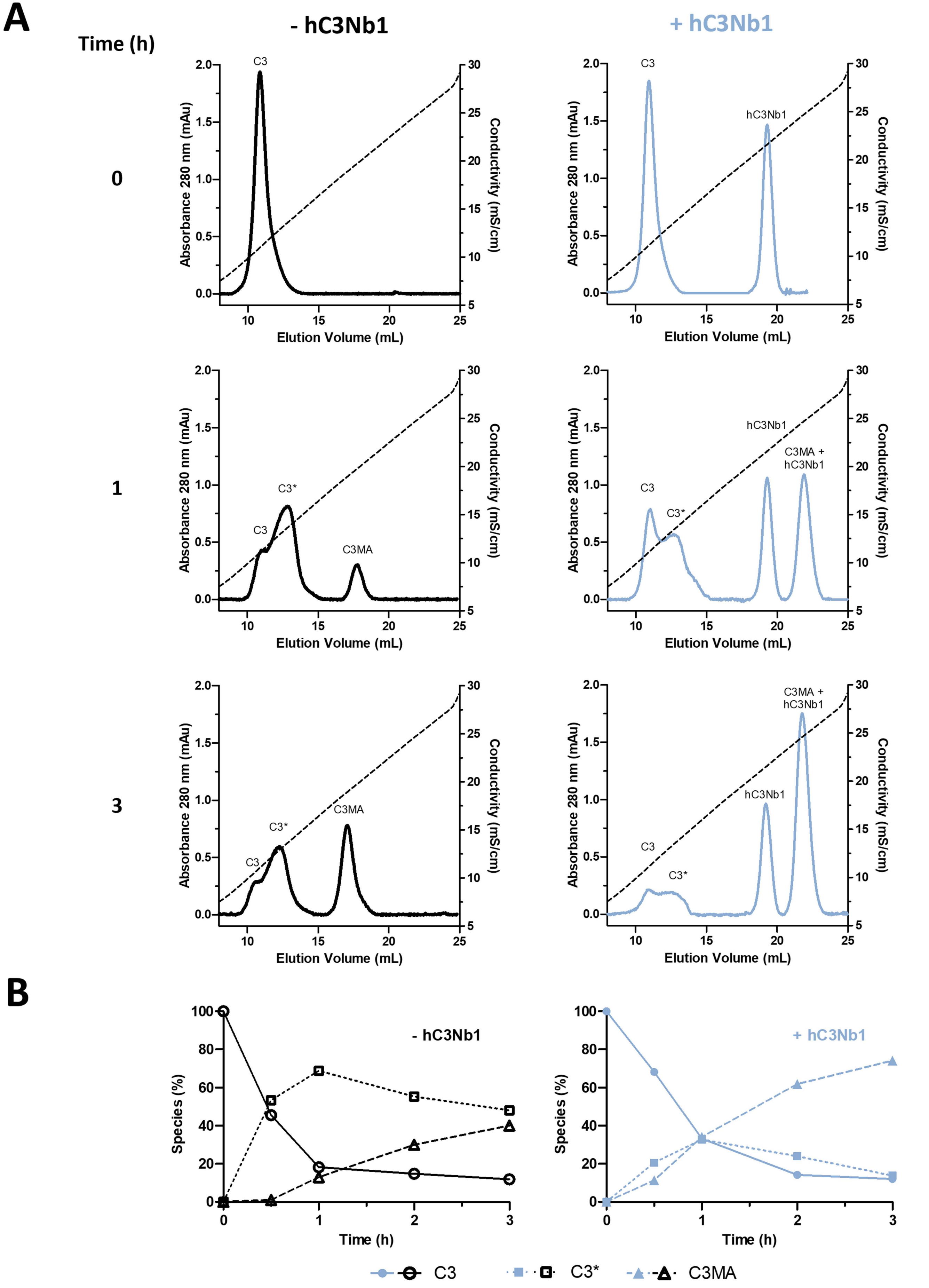
A
Time (h)
- hC3Nb1
+ hC3Nb1
0
1
3
C3
C3*
C3MA
hC3Nb1
C3MA + hC3Nb1
Absorbance 280 nm (mAu)
Conductivity (mS/cm)
Elution Volume (mL)
B
- hC3Nb1
+ hC3Nb1
Species (%)
Time (h)
C3
C3*
C3MA

| Start residue | Begin residue | Short domain name | Domain name | α-helix | β-strand | loop |
|---|---|---|---|---|---|---|
| 23 | 127 | MG1 | macroglobulin 1 | 0 | 38 | 67 |
| 128 | 228 | MG2 | macroglobulin 2 | 0 | 50 | 51 |
| 229 | 350 | MG3 | macroglobulin 3 | 6 | 64 | 52 |
| 350 | 449 | MG4 | macroglobulin 4 | 0 | 35 | 65 |
| 450 | 558 | MG5 | macroglobulin 5 | 3 | 66 | 40 |
| 559 | 599 | MG6N | macroglobulin 6 N-terminal | 0 | 21 | 20 |
| 600 | 671 | LNK | Linker region | 24 | 5 | 37 |
| 672 | 748 | ANA | anaphylatoxin | 44 | 0 | 30 |
| 749 | 769 | Nt-α' | N-terminal of C3b α-chain | 0 | 0 | 12 |
| 770 | 827 | MG6C | macroglobulin 6 C-terminal | 0 | 31 | 27 |
| 828 | 933 | MG7 | macroglobulin 7 | 0 | 59 | 47 |
| 934 | 985 | CUBN | complement C1r/C1s, Uegf, Bmp1 N-terminal | 0 | 20 | 32 |
| 986 | 1289 | TE | thioester | 189 | 0 | 115 |
| 1290 | 1355 | CUBC | complement C1r/C1s, Uegf, Bmp1 C-terminal | 0 | 28 | 38 |
| 1356 | 1494 | MG8 | macroglobulin 8 | 9 | 58 | 72 |
| 1495 | 1663 | C345c | C-terminal parts of netrins, complement proteins | 49 | 47 | 73 |

**Supporting table 1. Domains in human complement C3.** Notice that the MG6 and CUB domains contain inserts between their N-terminal and C-terminal parts. Numbers in α-helix, β-strand and loop columns were calculated in Pymol by counting the number of residues in complement C3 from pdb entry 6ru5 with secondary structure equal to “H”, “S” and “L”. Disordered residues 740-750 in the scissile bond region not modelled were assigned to “L” and shared between the ANA and Nt-α' rows.

| | C3MA-hC3Nb1 N108C dimer |
|---|---|
| **Data collection and processing** | |
| Microscope | Titan Krios G3i |
| Camera | Gatan K3 |
| Voltage (kV) | 300 |
| Magnification | 130,000x |
| Calibrated pixel size (Å) | 0.647 |
| Defocus range (µm) | 0.2-2.4 |
| Exposure on sample ($e^-$/Å$^2$) | 59.5 |
| Dose fractions | 53 |
| Exposure time (s) | 1.4 |
| Collected movies | 6,093 |
| Micrographs used for picking | 5,943 |
| Particle picks | 2,227,588 |
| Particles in final stack | 197,438 |
| Symmetry imposed | C1 |
| Global resolution at FSC = 0.143 (Å) | 2.9 |
| **Refinement** | |
| Initial model used (PDB code) | |
| C3MA | 6EHG |
| hC3Nb1 | 6EHG |
| Model resolution (Å) | 3.4 |
| FSC threshold | 0.5 |
| Map sharpening *B* factor (Å$^2$) | 59.2 |
| Model composition | |
| Atoms | 26958 |
| Protein residues | 3358 |
| Ligands | 2 BMA, 6 NAG |
| B factors (Å$^2$, min/max/mean) | |
| Protein | 43.9/441.2/177.2 |
| Ligand | 79.2/190.4/138.6 |
| R.m.s. deviations | |
| Bond lengths (Å) | 0.005 |
| Bond angles (°) | 1.044 |
| Validation | |
| MolProbity score | 1.62 |
| Clashscore | 4.22 |
| Poor rotamers (%) | 1.9 |
| CaBLAM outliers (%) | 1.68 |
| Ramachandran plot | |
| Favored (%) | 96.77 |
| Allowed (%) | 3.23 |
| Disallowed (%) | 0 |
| Rama-Z (RMSD) | |
| Whole (N=3346) | -1.4 (0.13) |
| Helix (N=524) | -2.42 (0.16) |
| Sheet (N=1312) | -0.05 (0.14) |
| Loop (N=1510) | -1.07 (0.14) |

**Supporting Table 2: Cryo-EM data collection, map and model refinement and validation statistics for the C3MA-hC3Nb1 dimer.**

| Label | Calculated mass (Da) | Modifications | Sequence |
|---|---|---|---|
| A0 | 6843.297 | None | MDKVGKYPKELRKCCEDGMRENPMRFSCQRRTRFISLGEACKKVFLDCCNYITELRR |
| A1 | 6859.292 | 1x Methionine oxidation | MDKVGKYPKELRKCCEDGMRENPMRFSCQRRTRFISLGEACKKVFLDCCNYITELRR |
| A2 | 6875.287 | 2x Methionine oxidation | MDKVGKYPKELRKCCEDGMRENPMRFSCQRRTRFISLGEACKKVFLDCCNYITELRR |
| A3 | 6891.281 | 3x Methionine oxidation | MDKVGKYPKELRKCCEDGMRENPMRFSCQRRTRFISLGEACKKVFLDCCNYITELRR |
| B0 | 7335.552 | None | MDKVGKYPKELRKCCEDGMRENPMRFSCQRRTRFISLGEACKKVFLDCCNYITELRRQHAR |
| B1 | 7351.547 | 1x Methionine oxidation | MDKVGKYPKELRKCCEDGMRENPMRFSCQRRTRFISLGEACKKVFLDCCNYITELRRQHAR |
| B2 | 7367.542 | 2x Methionine oxidation | MDKVGKYPKELRKCCEDGMRENPMRFSCQRRTRFISLGEACKKVFLDCCNYITELRRQHAR |
| B3 | 7383.537 | 3x Methionine oxidation | MDKVGKYPKELRKCCEDGMRENPMRFSCQRRTRFISLGEACKKVFLDCCNYITELRRQHAR |
| C0 | 7784.826 | None | SVQLTEKRMDKVGKYPKELRKCCEDGMRENPMRFSCQRRTRFISLGEACKKVFLDCCNYITELRR |
| C1 | 7800.821 | 1x Methionine oxidation | SVQLTEKRMDKVGKYPKELRKCCEDGMRENPMRFSCQRRTRFISLGEACKKVFLDCCNYITELRR |
| C2 | 7816.816 | 2x Methionine oxidation | SVQLTEKRMDKVGKYPKELRKCCEDGMRENPMRFSCQRRTRFISLGEACKKVFLDCCNYITELRR |
| C3 | 7832.811 | 3x Methionine oxidation | SVQLTEKRMDKVGKYPKELRKCCEDGMRENPMRFSCQRRTRFISLGEACKKVFLDCCNYITELRR |
| D0 | 8277.082 | None | SVQLTEKRMDKVGKYPKELRKCCEDGMRENPMRFSCQRRTRFISLGEACKKVFLDCCNYITELRRQHAR |
| D1 | 8293.077 | 1x Methionine oxidation | SVQLTEKRMDKVGKYPKELRKCCEDGMRENPMRFSCQRRTRFISLGEACKKVFLDCCNYITELRRQHAR |
| D2 | 8309.072 | 2x Methionine oxidation | SVQLTEKRMDKVGKYPKELRKCCEDGMRENPMRFSCQRRTRFISLGEACKKVFLDCCNYITELRRQHAR |
| D3 | 8325.067 | 3x Methionine oxidation | SVQLTEKRMDKVGKYPKELRKCCEDGMRENPMRFSCQRRTRFISLGEACKKVFLDCCNYITELRRQHAR |
| E0 | 9082.538 | None | SVQLTEKRMDKVGKYPKELRKCCEDGMRENPMRFSCQRRTRFISLGEACKKVFLDCCNYITELRRQHARASHLGLAR |
| E1 | 9098.533 | 1x Methionine oxidation | SVQLTEKRMDKVGKYPKELRKCCEDGMRENPMRFSCQRRTRFISLGEACKKVFLDCCNYITELRRQHARASHLGLAR |
| E2 | 9114.52752 | 2x Methionine oxidation | SVQLTEKRMDKVGKYPKELRKCCEDGMRENPMRFSCQRRTRFISLGEACKKVFLDCCNYITELRRQHARASHLGLAR |
| E3 | 9130.52243 | 3x Methionine oxidation | SVQLTEKRMDKVGKYPKELRKCCEDGMRENPMRFSCQRRTRFISLGEACKKVFLDCCNYITELRRQHARASHLGLAR |

**Supporting Table 3. Masses of peptides in the two C3a$_T$ samples determined by intact mass LC-MS analysis, as shown in supporting figure 4B-C**. The sequences of the peptides and corresponding number of methionine oxidation modifications assigned to each detected peptide mass are given. Note that the sequences were identified exclusively on the basis of MS1 mass. The peptide sequence designated “D” was the major sequence in C3a$_T$ from both C3 and C3MA.

| | C3* | | |
|---|---|---|---|
| **Data collection and processing** | | | |
| Microscope | | Titan Krios G3i | |
| Camera | | Gatan K3 | |
| Voltage (kV) | | 300 | |
| Magnification | | 130,000x | |
| Calibrated pixel size (Å) | | 0.647 | |
| Defocus range (µm) | | 0.7-2.4 | |
| Stage tilt (°) | 0 | | 25 |
| Exposure on sample ($e^{-}$/Å$^{2}$) | 59.5 | | 60.3 |
| Dose fractions | 53 | | 53 |
| Exposure time (s) | 1.4 | | 1.4 |
| Collected movies | 7,366 | | 897 |
| Micrographs used for picking | 6,687 | | 897 |
| Particle picks | 1,662,530 | | 494,075 |
| Particles in final stack | 49,852 | | 6,680 |
| Map focus | MG-ring | TE-CUB | Consensus |
| Global resolution at FSC = 0.143 (Å) | 5.7 | 7.8 | 8.8 |
| Local resolution at FSC = 0.5 (Å) 25th percentile/median/75th percentile | 5.8/6.3/7.1 | 7.0/7.5/8.0 | 10.5/11.4/12.8 |
| Map sphericity | 0.89 | 0.91 | 1.00 |

**Supporting table 4. CryoEM analysis C3*.** Data and map statistics for C3* image processing. Global resolution is given as GSFSC corrected by phase randomisation as reported by Non-Uniform/Homogenous Refinement in cryoSPARC. Local resolution distributions are given as reported by Local Resolution Estimation using a 20 voxel kernel.

| | C3*-hC3Nb2 | |
|---|---|---|
| **Data collection and processing** | | |
| Microscope | Titan Krios G3i | |
| Camera | Gatan K3 | |
| Voltage (kV) | 300 | |
| Magnification | 130,000x | |
| Calibrated pixel size (Å) | 0.647 | |
| Defocus range (µm) | 0.6-2.3 | |
| Stage tilt (°) | 0 | 20 |
| Exposure on sample ($e^-$/Å$^2$) | 59.3 | 59.7 |
| Dose fractions | 53 | 53 |
| Exposure time (s) | 1.4 | 1.4 |
| Collected movies | 1,412 | 582 |
| Micrographs used for picking | 1,053 | 368 |
| Particle picks | 142,134 | 87,353 |
| Particles in final stack | 14,700 | 7,350 |
| Global resolution at FSC = 0.143 (Å) | 4.2 | |
| Local resolution at FSC = 0.5 (Å) | | |
| 25th percentile/median/75th percentile | 4.6/5.1/6.2 | |
| Map sphericity | 0.59 | |

**Supporting table 5. CryoEM analysis C3* in complex with hC3Nb2.** Data and map statistics for C3*-hC3Nb2 image processing. Global resolution is given as GSFSC corrected by phase randomisation as reported by Non-Uniform in cryoSPARC. Local resolution distribution is given as reported by Local Resolution Estimation using a 31 voxel kernel.

| Domain 1 | Domain 2 | human C3 6ru5 | human C3 2a73 | bovine C3 8cem-1 | bovine C3 8cem-2 | C3b 6ehg | C3b 5fo7 | C3MA 8oq3-1 | C3MA 8oq3-2 | Average C3 | St dev C3 | Average C3b/C3MA | St dev C3b/C3MA | C3 - (C3b or C3MA) |
|---|---|---|---|---|---|---|---|---|---|---|---|---|---|---|
| TED | MG8 | 2464 | 1979.2 | 2546.2 | 2414.9 | -6 | -7.3 | -6.9 | -7.1 | 2351 | 254 | -7 | 1 | 2358 |
| CUB | TED | 2127.1 | 2214.2 | 2173.6 | 2158.3 | 681 | 553.9 | 697.7 | 596.1 | 2168 | 36 | 632 | 69 | 1536 |
| ANA | MG8 | 1491.8 | 1400.4 | 1304.7 | 1333.1 | 0 | 0 | 0 | 0 | 1383 | 83 | 0 | 0 | 1383 |
| MG2 | TED | 1088.6 | 1148.9 | 1035.6 | 1081.8 | -3.2 | -4.7 | -2.7 | -3.1 | 1089 | 47 | -3 | 1 | 1092 |
| MG3 | Nt-α' | 1219.6 | 1320.2 | 1047.7 | 1208.9 | 128 | 130 | 97 | 102.2 | 1199 | 113 | 114 | 17 | 1085 |
| ANA | Nt-α' | 969 | 922.6 | 566.5 | 689.8 | 0 | 0 | 0 | 0 | 787 | 191 | 0 | 0 | 787 |
| MG2 | Nt-α' | 695.9 | 771 | 988 | 1113.5 | 195.3 | 203.8 | 172.4 | 176.7 | 892 | 193 | 187 | 15 | 705 |
| CUB | MG8 | 1089.5 | 923.5 | 697.2 | 725.5 | 295.1 | 369.3 | 181.7 | 180.4 | 859 | 184 | 257 | 92 | 602 |
| MG3 | ANA | 689.7 | 634.6 | 497 | 578.1 | 0 | 0 | 0 | 0 | 600 | 82 | 0 | 0 | 600 |
| CUB | C345C | 898.9 | 676.5 | 713.2 | 748.7 | 184.5 | 217.5 | 156.1 | 201.5 | 759 | 98 | 190 | 26 | 569 |
| MG8 | C345C | 1053.6 | 1044 | 1206 | 1150.5 | 697.2 | 784 | 659.4 | 685.6 | 1114 | 78 | 707 | 54 | 407 |
| LNK | TED | -5.1 | -4.3 | -6.9 | -5.3 | 338.7 | 362.4 | 314 | 333.4 | -5 | 1 | 337 | 20 | -342 |
| MG3 | MG7 | 337.6 | 451.7 | 439.1 | 281.5 | 771.6 | 809.3 | 793.7 | 769.3 | 377 | 82 | 786 | 19 | -409 |
| MG2 | MG7 | 1.9 | 2 | 1.8 | 2.1 | 473.7 | 470.5 | 463.8 | 451.6 | 2 | 0 | 465 | 10 | -463 |
| MG3 | MG6 | 2.1 | 2.1 | 2 | 2.2 | 496.3 | 491 | 447.4 | 446.9 | 2 | 0 | 470 | 27 | -468 |
| MG3 | LNK | 2.2 | 2.1 | 2 | 2 | 530.3 | 579.8 | 433.9 | 445.1 | 2 | 0 | 497 | 70 | -495 |
| MG3 | MG8 | 484.1 | 434.2 | 473 | 545.6 | 1399.6 | 1516.4 | 1378.8 | 1425.5 | 484 | 46 | 1430 | 61 | -946 |
| MG2 | CUB | 2.3 | 2.5 | 2.1 | 2.5 | 1274.3 | 856.5 | 993.8 | 1107 | 2 | 0 | 1058 | 177 | -1056 |
| MG1 | TED | -6.2 | -5 | -7.2 | -5.1 | 1249.3 | 1174.3 | 1086.3 | 1222 | -6 | 1 | 1183 | 72 | -1189 |
| Nt-α' | MG7 | 0.4 | 0.4 | 0.4 | 0.5 | 1331.1 | 1597 | 1541.8 | 1453 | 0 | 0 | 1481 | 116 | -1481 |

**Supporting table 6. Buried surface areas in domain interactions.** For human C3, one molecule each from the pdb entries 2a72 and 6ru5 were analyzed while for bovine C3 two molecules from the entry 8cef were analyzed. For C3b, one molecule each from entries 6ehg and 5fo7 were analyzed while for C3MA, two molecules from entry 8oq3 were analyzed. The domains in human C3 were defined as in supporting table 1 and the corresponding domains in bovine C3 according to a pairwise alignment with human C3. Buried surface areas were calculated with PyMOL. Numerical differences less than 300 Å$^2$ are not shown.
